# Event-based Single Molecule Localization Microscopy (*eventSMLM*) for High Spatio-Temporal Super-resolution Imaging

**DOI:** 10.1101/2023.12.30.573392

**Authors:** Jigmi Basumatary, S Aravinth, Neeraj Pant, Vignesh Ramanathan, Chetan Singh Thakur, Partha Pratim Mondal

## Abstract

Photon emission by single molecules is a random event with a well-defined distribution. This calls for event-based detection in single-molecule localization microscopy. The detector has the advantage of providing a temporal change in photons and emission characteristics within a single blinking period (typically, ∼ 30 *ms*) of a single molecule. This information can be used to better localize single molecules within a user-defined collection time (shorter than average blinking time) of the event detector. The events collected over every short interval of time / collection time (∼ 3 *ms*) give rise to several independent temporal photon distributions (*tPSFs*) of a single molecule. The experiment showed that single molecules intermittently emit photons. So, capturing events over a shorter period / collection time than the entire blinking period gives rise to several realizations of the temporal PSFs (*tPSFs*) of a single molecule. Specifically, this translates to a sparse collection of active pixels per frame on the detector chip (image plane). Ideally, multiple realizations of single-molecule *tPSF* give several position estimates of the single-molecules, leading to multiple *tPSF* centroids. Fitting these centroid points by a circle provides an approximate position (circle center) and geometric localization precision (determined by the FWHM of the Gaussian) of a single molecule. Since the single-molecule estimate (position and localization precision) is directly driven by the data (photon detection events on the detector pixels) and the recorded *tPSF*, the estimated value is purely experimental rather than theoretical (Thomson’s formula). Moreover, the temporal nature of the event camera and *tPSF* substantially reduces noise and background in a low-noise environment. The method is tested on three different test samples (1) Scattered Cy3 dye molecules on a coverslip, (2) Mitochondrial network in a cell, and (3) Dendra2HA transfected live NIH3T3 cells (Influenza-A model). A super-resolution map is constructed and analyzed based on the detection of events (temporal change in the number of photons). Experimental results on transfected NIH3T3 cells show a localization precision of ∼ 10 *nm*, which is ∼ 6 fold better than standard SMLM. Moreover, imaging HA clustering in a cellular environment reveals a spatio-temporal PArticle Resolution (PAR) (2.3*l*_*p*_ × *τ*) of 14.11 *par* where 1 *par* = 10^−11^ *meter*.*second*. However, brighter probes (such as Cy3) are capable of ∼ 3.16 *par*. Cluster analysis of HA molecules shows > 81% colocalization with standard SMLM, indicating the consistency of the proposed *eventSMLM* technique. The single-molecule imaging on live cells reveals temporal dynamics (migration, association, and dissociation) of HA clusters for the first time over 60 minutes. With the availability of event-based detection and high temporal resolution, we envision the emergence of a new kind of microscopy that is capable of high spatio-temporal particle resolution in the sub-10 *par* regime.

## I. INTRODUCTION

With the arrival of super-resolution microscopy, it is now possible to unveil the nanoscopic world of single molecules and their interaction in a cellular system. The techniques that largely facilitate super-resolution are either based on structured illumination or stimulated emission depletion (STED) or single molecule blinking (such as PALM, fPALM, STORM). These techniques have advanced so much that they are able to resolve features down to a few tens of nanometers. Needless to say, these techniques hold the potential to reveal biological processes in their natural environment.

Early 1990s saw the emergence of a new concept (called STED) that for the first time demonstrated the capability to surpass the diffraction limit [7] [1] [8]. It wasn’t until early 2000s that the first far-field super-resolution technique is fully established [8]. Then the year 2005-2006 saw the emergence of two different techniques (structured illumination and single molecule localization microscopy) that showed resolution far below the classical diffraction limit [9] [3] [2] [4]. This established the fact that it is possible to surpass the diffraction limit that was once thought impossible. Subsequent years saw the success of super-resolution technique and emergence of a family of important variants. Most of these variants were inspired by applications in diverse disciplines ranging from biological to physical sciences. This include the development of STED [1] followed by stochastic optical reconstruction microscopy(STORM)[2], fluorescence photoactivated localization microscopy (f/PALM)[3][4] and DNApoint accumulation in nanoscale topography (PAINT)[5]. While STED is purely a RESOLFT based technique, PALM / fPALM / STORM are truly single molecule based techniques collectively known as Single Molecule Localization Microscopy (SMLM) [51]. Since the introduction of SMLM techniques, several variants are proposed to improve spatial resolution [10][11] [12][14][49][15]. Specifically, the detection of fortunate molecules (molecules that are bright and blink for long duration) has shown a significant PArticle Resolution Shift (PAR-shift) in both fixed and live cell specimens [16] [17]. Recent advances has seen emergence of a new techniques (MINFLUX and MINSTED) that combines both STED and single molecule localization. These techniques have demonstrated a spatial resolution in < 10 *nm* regime [18] [19]. Moreover, they have used single molecule properties such as, emitter orientation[27], molecule lifetime[31][32][33], and even information related to local neighborhood in a cellular system [28][29] to facilitate single-molecule orientation-localization microscopy [30], multicolor imaging [34][35] and single molecule tracking [45]. Although these super-resolution techniques can achieve spatial resolution in the range of 15−30 *nm*, but these resolutions do not translate directly to live cell experiments. This becomes even more challenging for observing single molecules and its collective dynamics due to poor temporal resolution.

With the emergence of new applications, the need for high spatio-temporal resolution has gained interest. This calls for better localization precision and high temporal resolution. To overcome these two key obstacles, we propose event based detection of single molecules, practically realized by event detector/camera. These specialized detectors are different from sCMOS / EMCCD in the sense they are based on differential intensity, which means that event camera is sensitive to pixel-level brightness / intensity changes (Δ*I*) instead of intensity (*I*) itself. They offer many advantages over traditional detectors (CCD EMCCD sCMOS) such as, near-zero motion-blur, high temporal resolution (in the order of few tens of microseconds), and negligible readout time. For standard frame-based detectors, each pixel synchronously integrates the incoming photon in a frame. This nature limits the information of sporadic fluorescence emission of scholastically activated single molecules. Recently developed bio-inspired event-based detector has demonstrated the highest reported temporal resolution (100, 000 *frames/sec*). This is due to the data-sensing architecture which is designed to monitor continuous changes in brightness (log intensity). When the changes in magnitude exceed the predefined threshold magnitude at a particular time, the camera sends an event having pixel location (*x, y*), time (*t*), and the polarity (*p*) of the event. The polarity of an event is either positive (ON event) or negative (OFF event), corresponding to increased or decreased intensity, respectively. Hence event camera output is a variable sequence of events (ON/OFF) with time and polarity. Asynchronously detected events are transferred to the periphery processing unit using a fast readout bus called address-event representation (AER) readout bus and the events are recorded [36]. Detailed description of event cameras and its architecture can be found in Ref. [37] [38].

The use of event camera in SMLM can be advantageous, primarily due to event-based detection and high temporal resolution. These properties are exploited in a range of imaging modalities. In ion imaging, event-based camera is used to study real-time velocity-resolved kinetics of thermal desorption [40]. Recently event based detectors with deep learning approaches were used to localize and tract fluorescent particles below diffraction limit (< 50 *nm*) and with high temporal resolution (in milliseconds) [45]. Gu et al has demonstrated a single molecule detection method primarily based on the temporal pixel intensity fluctuation for rapid determination of localization of events with good sensitivity [39]. In another development, event-based detectors were used to image biological samples with a spatial resolution comparable to that of standard SMLM [42]. Of-late, event detectors have found applications in light field microscopy (LFM) where it has demonstrated reconstruction of fast-moving biological objects at kHz frame rates. Moreover, the technique (eventLFM) allowed imaging of blinking neuronal signal in brain tissue and tracking of GFP-labelled neurons in a moving C. elegans. This suggests that event based detection has the potential to record events faster and more reliably in SMLM.

The asynchronous recording of stochastic emission from single molecules in time is ideal for better determination of the position and localization of single molecules. The advantage of high temporal resolution and event sensitive detection allows recording of multiple realizations of the PSF within the blinking period of a single molecule. This has the advantage of recording fortunate molecules (molecules that are bright) for extended period of time, giving multiple PSF realizations. Ideally, centroids can be computed from individual *tPSFs* and mean can be calculated along with its localization geometrically.

The process largely bypasses background noise and noise pixelation factor as requited by standard SMLM. The technique is tested on multiple specimens such as, scattered Cy3 single molecules, and organelle-specific fluorophores in a fixed cell. In a separate experiment, a Influenza (type A) disease model is considered where a key glycoprotein Hemaglutinin (HA) is monitored in live NIH3T3 cell determining temporal dynamics of HA clusters. The high spatio-temporal resolution of event based detection technique is expected to substantially expand the reach of localization microscopy.

## II. RESULTS

The event detector used in *eventSMLM* microscopy enables temporal detection of photons (emitted by single molecule) during the ON time. This is very different from detectors generally used in SMLM where the detection of single molecules are carried by a frame based intensity CCD/EMCCD/sCMOS camera. These detectors work by recording integrated intensity at each pixel during ON time. Here we discuss event based detection carried out by an event detector.

### A. Event detection by Event Camera/Detector

As compared to the standard frame-based cameras (CCD/EMCCD/sCMOS) that record absolute brightness, event-based cameras respond to to brightness change asynchronously and independently for every pixel. Thus the output of an event-based detector are events, with each event correspond to a change of brightness (log intensity). Each pixel memorizes the log intensity at each time and continously monitors for a change. The camera senses an event for a change exceeding the threshold, upon which the location (*x, y*), the time (*t*) and the polarity (*p*) is recorded. Here the 1-bit polarity represents increased brightness (+1) or decreased brightness (−1). As far as the speed is concerned, an event camera can achieve readout rates of upto 2 MHz [44].

The pixels in an event camera are designed to record and respond independently. The pixels respond to changes in their photo-currents, *I*_*pc*_ = *log*(*I*) which correspond to brightness. An event *e*_*k*_ = (*r*_*k*_, *t*_*k*_, *p*_*k*_) triggered at a pixel *r*_*k*_ = (*x*_*k*_, *y*_*k*_) (at location, *r*_*k*_) at time *t*_*k*_ when the change in brightness since the last event, Δ*I*_*pc*_(*r*_*k*_, *t*_*k*_) = *I*_*pc*_(*r*_*k*_, *t*_*k*_) − *I*_*pc*_(*r*_*k*_, *t*_*k*_ − Δ*t*_*k*_) reaches a temporal contrast threshold (say, ±*T*, with *T* > 0) i.e,

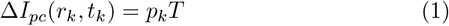

where, *p*_*k*_ = +1, − 1 is the polarity indicating increasing and decreasing brightness. Note that, Δ*t*_*k*_ is the time elapsed since the last event at the pixel, *r*_*k*_, and the pixel bias current determines the contrast sensitivity *T*. Typical thresholds are set between 15−50%, where the lower limit on *T* is determined by noise (short noise, thermal noise and circuit noise).

Note that an event camera sets a threshold on the change in brightness since the last event. An event is described as a change in brightness with time and can be interpreted as temporal derivative of brightness. For small Δ*t*_*k*_, the increment, Δ*I*_*pc*_(*r*_*k*_, *t*_*k*_) can be approximated by Taylor expansion. Incorporating the same in eqn(1) produces the following expression for an event,

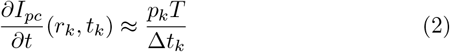

where, 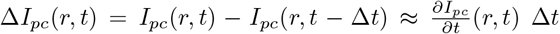 (Taylor expansion). A detailed description of an event detection can be found in **Supplementary 1**.

### B. Single Molecule Detection

A simple optical setup for event based detection is shown in Fig. 1A. The system is equipped with dual detectors i.e, standard EMCCD camera (Andor iXon Ultra 897) and an event camera (Prophesee, *EV KV* 2_*gen*_4.1, France). A 50:50 beam-splitter is introduced to split the expanded beam equally, which are then focused on to the cameras. This setup is to calibrate and evaluate the response of both the cameras under constant switch ON/OFF of photon stream from the light source (laser). Other optical components used to change the incident intensity are not shown. An electronic chopper is used to control ON / OFF state of the illumination light source (532 nm CW laser). Corresponding time diagram is shown in Fig. 1B. The EMCCD images are acquired during ON / OFF state of the laser and the data is obtained at a time interval of 50 *ms* (0−*T*) as shown in Fig. 1B. Fig. 1C shows the images captured by the event camera during the transition (*ON* →*OFF* / *OFF*→ *ON*) implying change in the number of photons and identify them as events. Positive and negative events corresponding to the polarity i.e, positive and negative change in intensity (red and blue) are also indicated along with the images. On the other hand, frame-based EMCCD camera fail to show the changes in the intensity during transition and in-fact records integrated intensity during the collection time. It is quite evident that event camera records non-zero images only during the rise and fall time and the associated polarity.

**FIG. 1:**
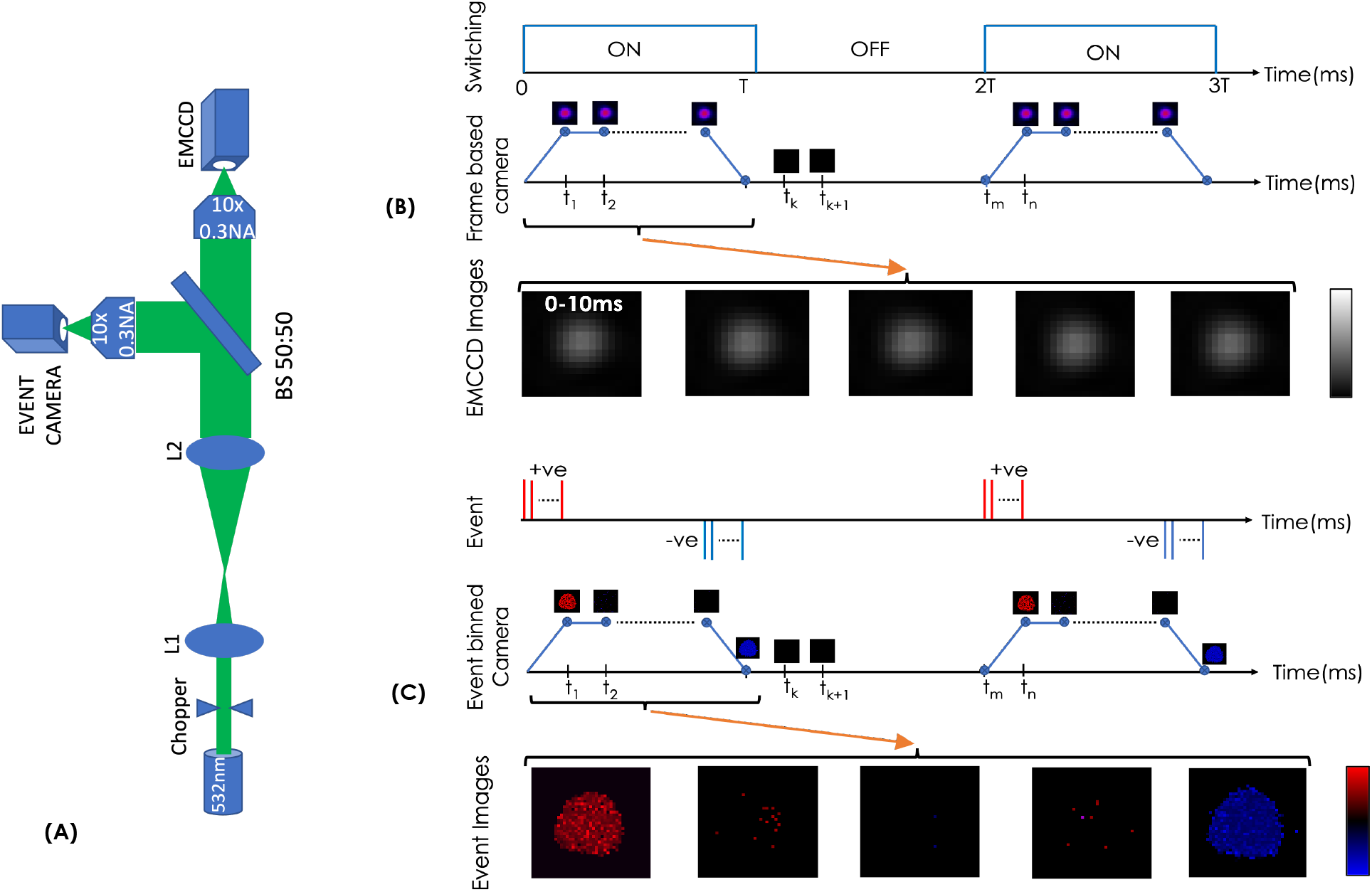
A. Light switch-on / off event of 532nm continuous laser. The beam chopper is motorized in which driving voltage applied to it is controlled via a relay. Frequency of relay switching is programmed using Arduino. More details can be found in **Supplementary Note 2**. During the ON time, chopper is open, so laser light continuously falls on both the detector, whereas chopper stops the laser during OFF time. [B,C] Timing diagram of ON/OFF events and corresponding data recorded by CCD (EMCCD with zero gain) and event camera. The relay switching is providing a means to synchronize data from both the cameras. However this needs to be taken care of during processing the recordings. During the ON time of the chopper CCD detector shows the Gaussian profile of laser spot. However in case of event detector series of events are observed at a pixel(*x, y*), only whenever there is a rising edge (red spikes) or falling edge(blue spikes) of ON/OFF. This shown that event detector records only whenever there is change in intensity. Red spikes represents positive events corresponding to rising edge whereas blue spike represents negative events corresponding to falling edge. The event-based frames are recorded over the desired collection times (here, 10 *ms*) by collecting events (corresponding to pixel intensity).

To characterise emission from a single molecule, we used event (temporal change in the number of photons) based detection scheme. The data is recorded from sparsely spread *Cy*3 single molecules on a coverslip at sampling rates of, 333 *Hz* (event collection time of *τ* = 3*ms*), 100 *Hz* (*τ* = 10*ms*) and 33 *Hz* (*τ* = 30*ms*). Fig. 2A shows the optical setup for simultaneous detection of single *Cy*3 molecules in both the detectors (EMCCD and Event cameras). The schematic diagram of detailed optical setup is shown in **Supplementary Note 3**).

**FIG. 2:**
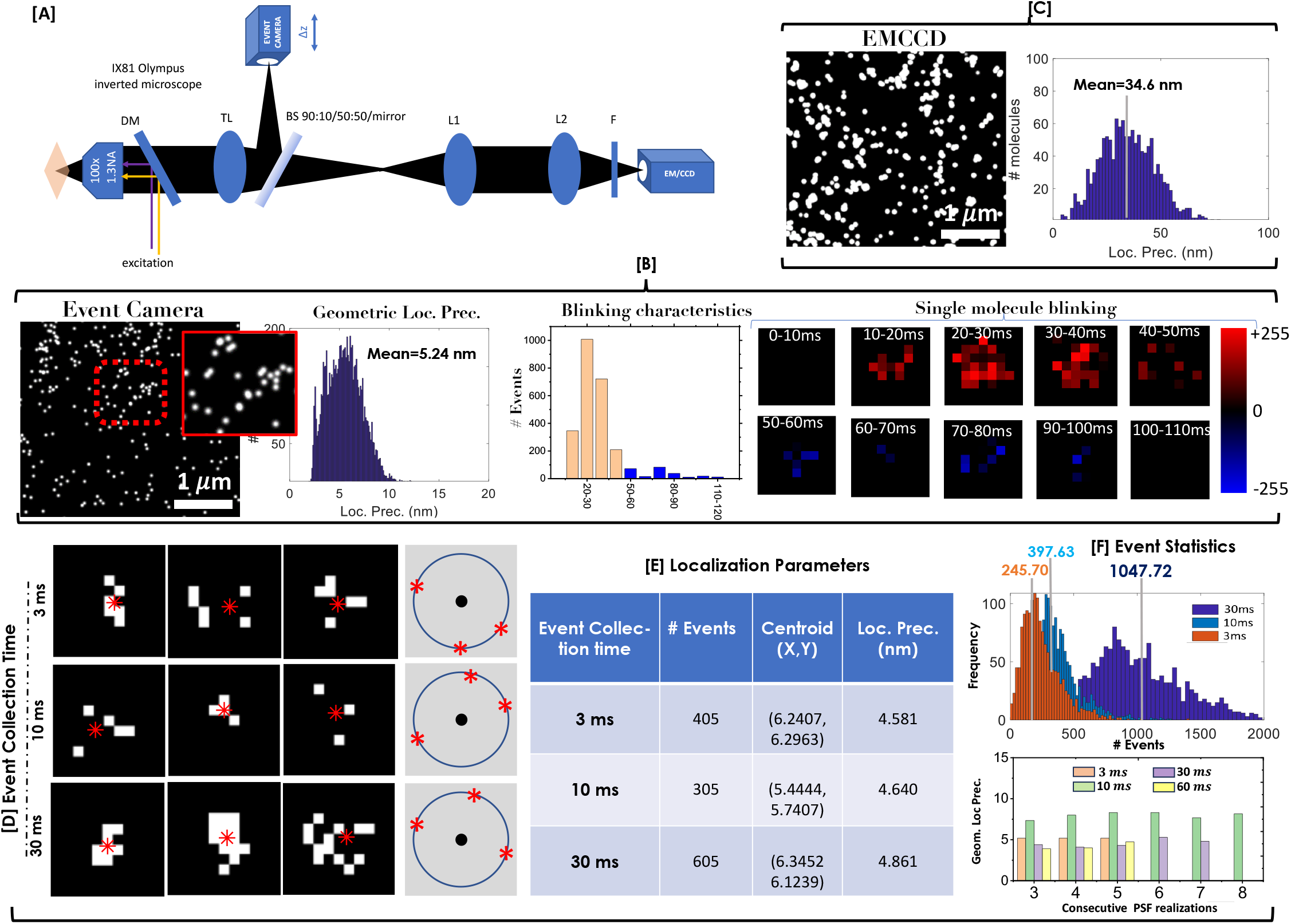
[A] Schematic diagram of event camera integration in the detection subsystem of conventional SMLM. The complete system is shown in **Supplementary Note 3**. The detection arm is designed for simultaneous two-channel imaging, with one arm for even-based detection using event camera and the other arm for single molecule detection using EMCCD camera. Event camera records the reflected light from the 50 : 50 beam splitter (BS), whereas EMCCD records transmitted light. A flipper mirror (not shown) is used in place of BS to image 100% signal form the event camera. [B] Event(both positive and negative) based reconstruction and geometric localization of reconstruction of cy3 molecules. Corresponding localization precision of 4.581 *nm* (average at an event collection time *τ* = 3*ms*) along with positive / negative event distribution frames are also shown. [C] EMCCD reconstructed recorded super resolved image using frame-based SMLM technique. Single molecules are localized at 34.6 *nm*. [D] Event based segmented *tPSF* (from both positive and negative events) with centroid at collection time of 3ms,10ms and 30ms collection time. A circle is fitted through centroid of the segmented *tPSFs*. Center and radius of the circles are determined and a Gaussian is associated from which position (centroid) and localization precision (FWHM) of the molecule are computed.[E] Centroid and geometric localization of three different single molecule *tPSFs*. [F] Distribution of number of event detected(both positive and negative) in Cy3 molecules at varying collection time of 3ms, 10ms and 30ms.

The event based reconstructed super-resolved image of Cy3 molecules is shown Fig. 2B. Localization precision of these molecules are calculated using geometric localization method (see, **Supplementary Note 4**). For comparison, reconstruction for standard SMLM (data acquired by EMCCD) along with localization precision (using Thompson’s formula) is also shown in Fig. 2C [50]. Fig. 2D shows the distribution of a typical single molecule captured by event camera at a time sampling of 3 *ms*, 10 *ms* and 30 *ms*. Since the molecule blinking period is much longer (in tens of milliseconds), the event camera captures multiple realizations of the bright PSF spots of a blinking molecule. From the recorded data/images it is apparent that the photon emission from single molecule is random within the blinking period, and the corresponding PSFs have distinct distributions. This is important from the point-of-view of super-resolution imaging, since this allows the localization of single molecules purely based on geometric parameters estimated from the recorded *tPSF*. Subsequently, the centroid of spot is determined from consecutive frames (see, red star in Fig. 2D). Assuming symmetric localization precision, a circle is fit (with a goodness-of-fit value, 0.9) to the centroids (blue circle) and the centre of circle (black dot) is determined (see Fig. 2D and IV.F). Subsequently, a Gaussian corresponding to the circle is identified and FWHM is calculated. This gives the position and localization precision (FWHM) for the single molecule. The corresponding localization parameters for a typical single molecule is tabulated in Fig. 2E, and detected event statistics at varying collection times is shown in Fig. 2F. The same process is repeated for all the molecules and a super-resolved map is constructed. The details of calculation for geometric localization precision is discussed in **Supplementary Note 4**. The localization precision plots indicate an average localization of 4.581 *nm* at *τ* = 3 *ms* of a typical *Cy*3 molecule. This is about 7 times less than the localization precision of standard SMLM that use well-known Thompson’s formula [50]. The corresponding spatial resolution is approximately. *R* = 2.3*l*_*p*_, where *l*_*p*_ is the localization precision [51]. Overall, this suggests a spatio-temporal particle resolution of 2.3*l*_*p*_ × *τ* = 2.3×4.581×3 *nm*.*msec* = 3.16 *par*. This is way better than standard SMLM.

### C. Imaging Mitochondrial Network in a Cell

To substantiate the new technique, we imaged mitochondrial membrane protein (Tom20) labelled with mEos photoactivable probe. Fig. 3A shows the epifluorescence image of the mEos-Tom20 transfected image of which a small region is chosen for study. The corresponding event based super-resolved image is shown in Fig. 3B. Alongside, the reconstructed SMLM image (reconstructed from the data/images captured by EMCCD camera) is also displayed for comparison (see, Fig. 3C). Three sub-regions (R1, R2 and R3) are chosen and compared as shown in Fig. 3(B,C). The corresponding localization plot gives an average localization precision of 5.677 *nm* for *eventSMLM* reconstructed image. This is way below the localization precision of standard SMLM which is 40.03 *nm*. Note that the data is obtained at *τ* = 10 *ms*. Super-resolved images at other event collection times (10 − 100 *ms*) are shown in Fig. S5-1 **Supplementary Note 5**.

**FIG. 3:**
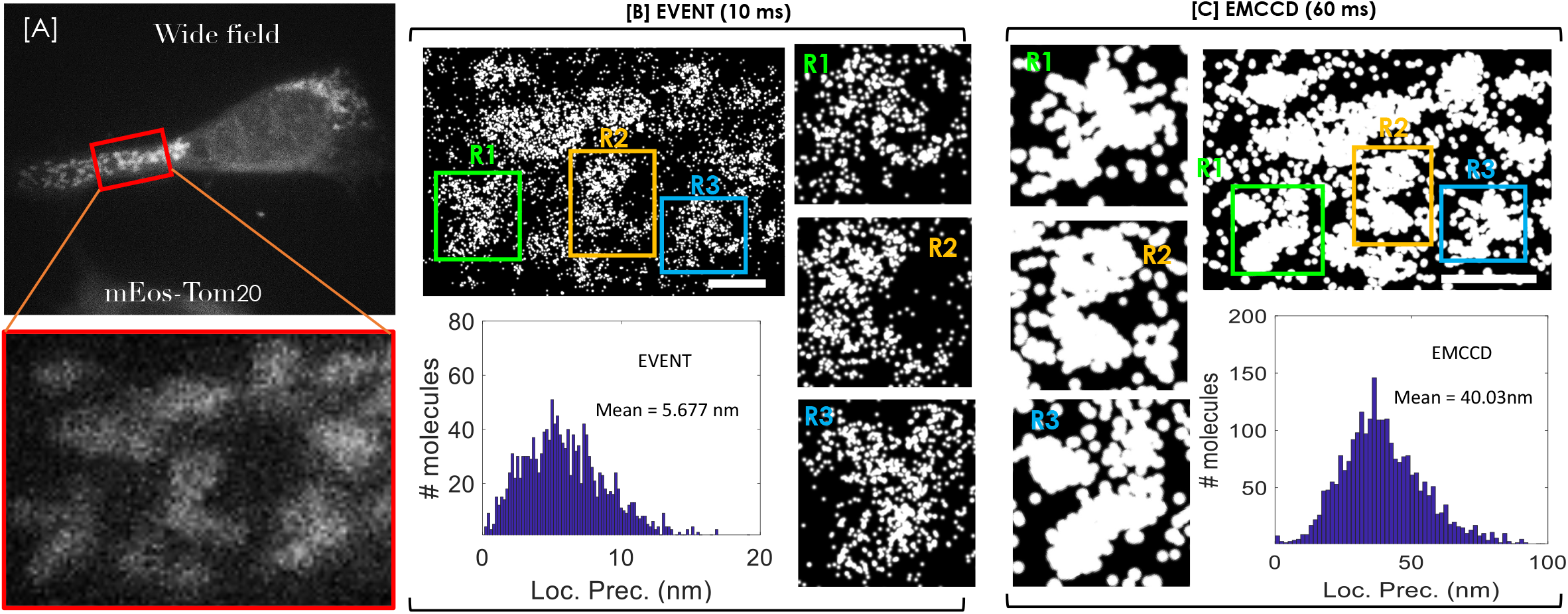
[A] Widefield fluorescence image of mEosTom20 transfected fixed NIH3T3 cell. Red doted zone is ROI. [B] Event detection based reconstructed superresolved image of Tom20 molecules. Frames are constructed by collecting events over 10ms. In the frames each pixel gray levels equal to the sum of negative and positive events detected in 10ms. [C] Localization map of Tom20 molecules detected using EMCCD (30ms exposure time). Enlarge views show of regions R1, R2 and R3 from [B] and [C] along with respective localization precision histogram. Data presented is simultaneous imaging using event and EMCCD camera.

### D. Imaging HA Clusters in a Cell

We employ *eventSMLM* system to understand spatio-temporal kinetics of HA molecules and its collective dynamics in an Influenza (type-A) disease model. Healthy NIH3T3 cells were transfected with a viral plasmid DNA (Dendra2-HA). This is followed by incubation at a temperature of 37°*C* and 5% supply of *CO*_2_. One set of cells were fixed by standard protocol (see, Methods section, IV.H) and the other set is washed with PBS and prepared for live imaging. Fig. 4A shows widefield fluorescence image of a fixed cell. The corresponding reconstruction using event camera is shown in Fig. 4B along with SMLM reconstructed image (Fig. 4C.) Note that, SMLM super-resolved image is reconstructed from single molecule data collected using EMCCD camera. Specifically, two regions (R1 and R2) were chosen for detailed study as displayed in Fig. 4(B,C). It is visually evident that clusters are better resolved in *eventSMLM* reconstructed image. This is due to high localization of single molecules, in-fact sub-clusters begin to appear within large clusters. Event based super-resolved images of HA clusters at other collection time (10 − 100 *ms*) is shown in **Supplementary Note 6**. Alongside, the localization study is also shown which indicate an average localization precision of 6.138 *nm*. Since the frame is captured at 10 *ms* (with 100*µsec* read-time for event camera) as compared to 60 *ms* (30 *ms* for capture + 30 *ms* for read/write) for SMLM, this shows ∼6 fold improvement in data acquisition per frame. Overall the system suggests an impressive spatio-temporal resolution (2.3*l*_*p*_ × *t*) of ∼2.3 × 6.138 × 10 *nm. msec* = 14.11 *par*, where, 1 *par* = 10^−11^ *meter*.*second*. This is approximately, 17.84 times better than SMLM for which standard spatio-temporal resolution is, ∼2.3 × 17.84 × 30 *par*. The corresponding FRC analysis for both the samples (mEos-Tom20 and Dendra2-HA) is discussed in **Supplementary Note 7**.

**FIG. 4:**
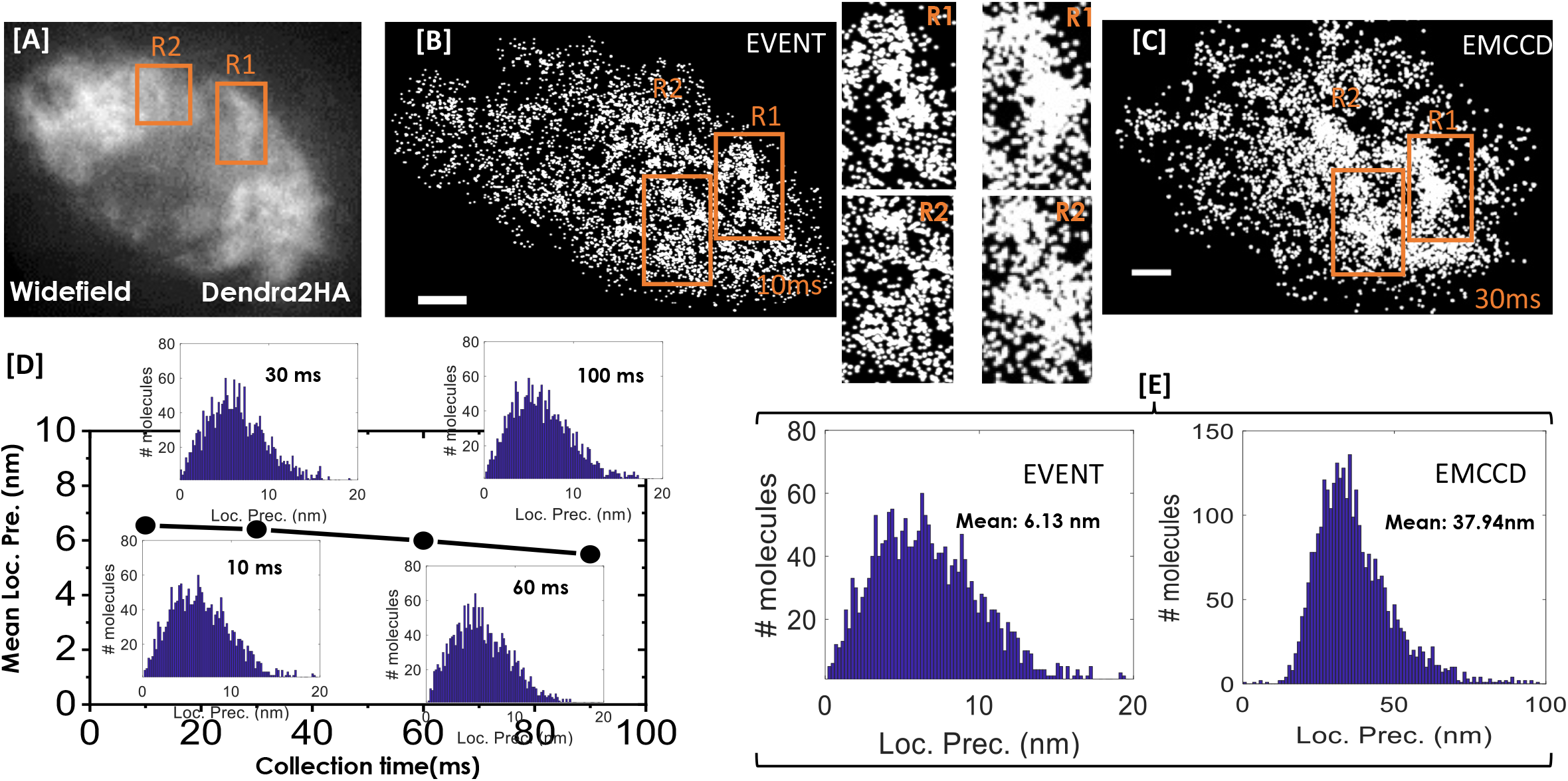
[A] Widefield fluorescence image of Dendra2HA transfected fixed NIH3T3 cell. [B] Event based reconstructed superresolved image. Frames are constructed by collecting all the events over 10ms. [C] Super-resolved map (SMLM) of HA molecules recorded by EMCCD (30ms exposure time). Enlarge view of ROIs (R1 and R2) also shown. [D] Event based Localization precision variation with event collection time. [E] Localization precision of the molecules detected by event detector(left) and EMCCD (right). Data presented is simultaneously detected by event and EMCCD using a 90:10 BS. Scale bar = *μ*m.

To further validate the potential of event-based single molecule super-resolution microscopy (*eventSMLM*), we have used it to visualize HA single molecules in Dendra2-HA transfected NIH3T3 cells (The cells were fixed post 24 hrs of transfection). The reconstructed super-resolved image of HA distribution (only clustered molecules) at varying event collection time is shown in Fig. 5A. Alongside overlay of all the images are displayed that indicate more HA molecules per cluster for 100 *ms*, thereby completing the finer features of individual clusters. In addition, a comparison of event based detection (data acquired using event camera) and frame based detection (EMCCD camera used for SMLM) is shown in Fig. 5B. Note that, simultaneous data acquisition is achieved for both the camera using 90 : 10 beamsplitter (90% for event camera and 10% for EMCCD) for comparison. Alongside an overlay of both the data suggests good colocalization of 0.8156. In addition, we have estimated biophysical parameters independently for both the detection schemes (event and frame) as shown in Table S9-T (see, **Supplementary Note 9**). A strong overlap of clusters indicate reliability of event-based super-resolution microscopy (see, Fig. 5B). This strengthens the fact that, event-based detection has high degree of reliability along with the added advantage of high spatio-temporal resolution.

**FIG. 5:**
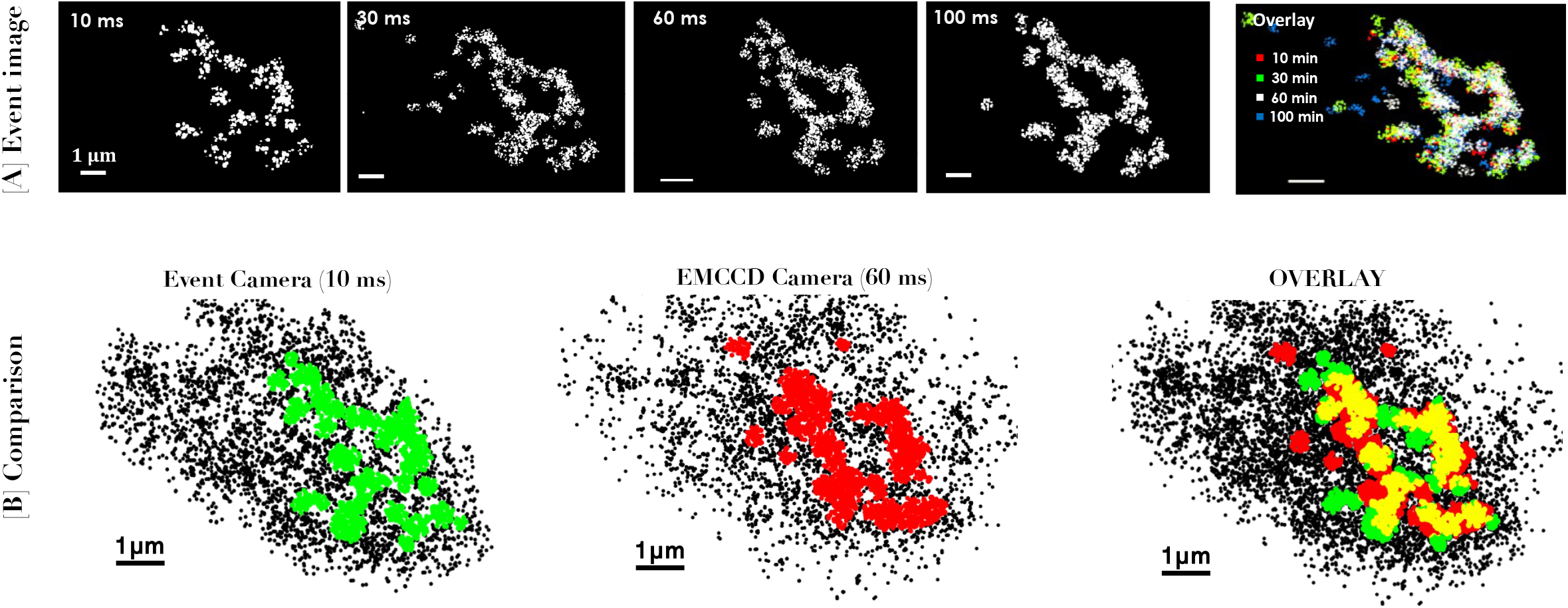
[A] DBSCAN algorithm applied to identify the cluster of HA molecules in a fixed NIH3T3 cell at varying collection event time of 10ms, 30ms, 60 ms and 100ms. [B] Event detected HA cluster (10ms event collection time) is compared with EMCCD detected cluster. The cluster is found to be consistent(see overlay image) with colocalization factor of 0.8156.

### E. Time-lapse Imaging of Dynamic HA Clusters

To access the potential of *eventSMLM* system, we have used it to visualize HA dynamics in live cells. The system is configured to take time-lapse data / images at a rate of 10 *ms* after every 15 minutes. Note that, event camera take continuous data with a negligible registration time (order of few tens of microseconds) [37] [38], whereas EMCCD camera needs about 30 *msec* for data registration after collecting data for each frame. This makes event camera as an ideal detector for imaging dynamical events. For the live experiment, the data is collected continuously over 60 minutes by the event detector, and the same is shown in Fig. 6A. Prominent clusters are identified using DBSCAN algorithm [53] and the individual clusters are colored (see, Fig. 6A). In addition, a separate binary cluster map consists of only the clustered molecules is shown in Fig. 6B. Alongside, the overlay of all the images are also displayed that indicates dynamic evolution, formation and even migration of HA clusters in the live cell. The corresponding plot show negligible change in localization precision for data taken over 60 minutes suggesting the bias-free data acquisition without significant photo-bleaching effect (see, Fig. 6C). We have also estimated biophysical parameters at every time-point as shown in Fig. 6(D,E). This indicates increase in cluster area and number of HA molecules per cluster and an increased cluster fraction suggesting increasing HA participating in cluster formation post 24 hrs of transfection. More details related to HA cluster dynamics is discussed in **Supplementary Note 10**.

**FIG. 6:**
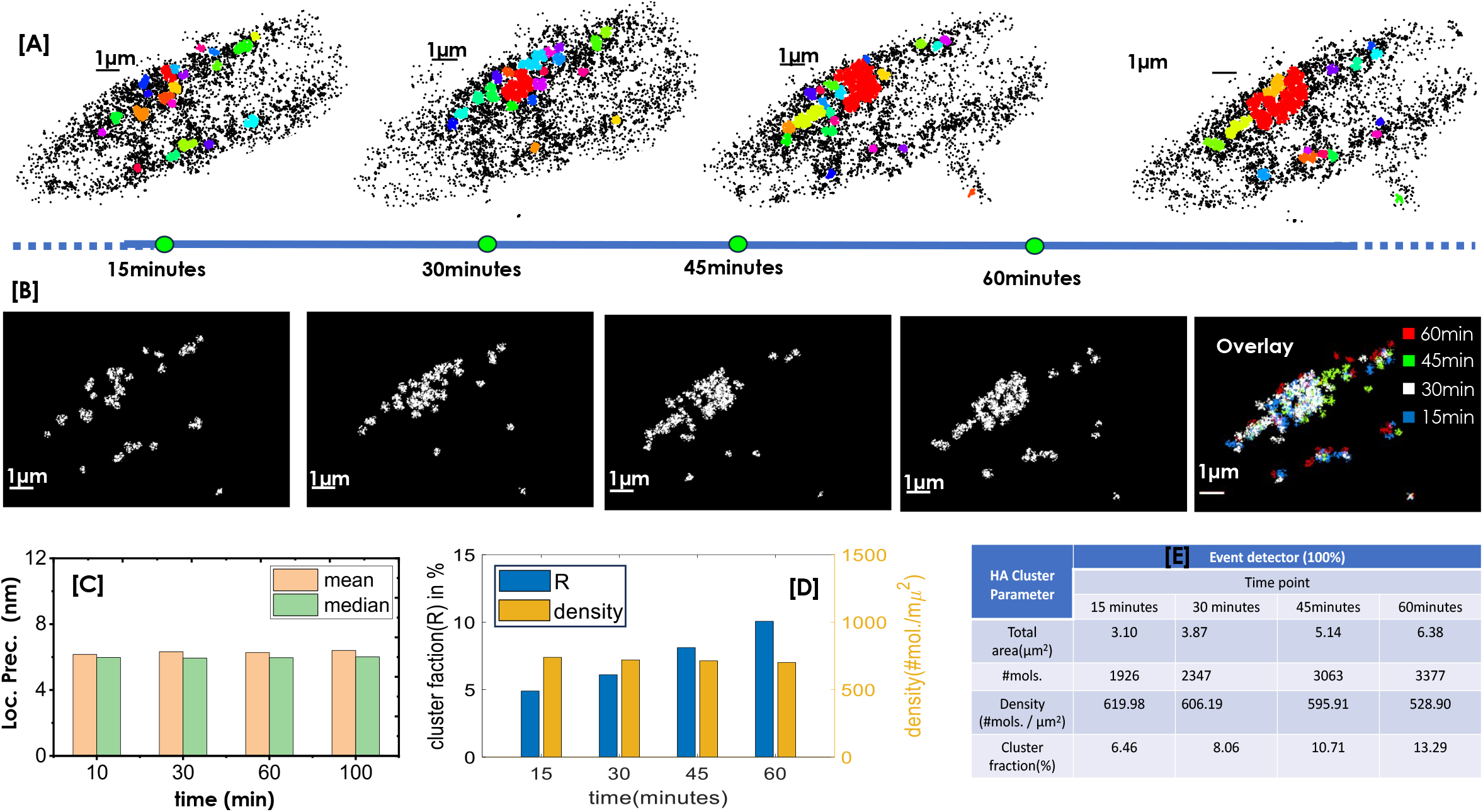
Live Cell Imaging: [A] DBSCAN algorithm applied to identified the dynamics of HA clusters in *eventSMLM* super-resolve images. Frames were reconstructed using 10 ms event collection time. [B] Map of only clustered HA molecules (unclustered molecules are removed). Color coded time overlay are also shown. [C] Variation of localization precision with time. [D] Change in cluster fraction (percentage of clustered molecules) and density of the HA molecules in clusters with time. [E] Tabulated biophysical parameters related to HA clusters. Mirror is used to detect 100% signal / fluorescence.

The high spatio-temporal resolution of *eventSMLM* allows dynamic tracking of HA clusters and their ensemble behavior in a live cell. To facilitate time-lapse study, raw images are recorded and processed to reconstruct the single molecule map at each time-point. We have chosen a 15 *min* time-scale to look for changes in the HA clusters for the time-lapse imaging. To facilitate single molecule analysis, we have used DBSCAN clustering method [55]. Since our interest lies in the study of HA clustering process, non-clustered molecules are removed and only the molecule involved in clustering are retained. Fig. 7A shows the time-lapse super-resolved images of HA clusters post 24 hrs of transfection. The clusters are color coded to represent the dynamic changes at single molecule level. The corresponding color code is represented below. Accordingly, all the stationary clustered are marked white. The increase in the number of molecules and overall size of cluster is indicated by single (↑), double (↑↑) and triple (↑↑↑) arrows pointing-up, whereas the decline in the number of molecules in a cluster is shown by arrows pointing down. The increase in subsequent frames are represented by, orange, red and brown, whereas, decrease is represented by dark-blue and light-blue. Apart from standard increase and decrease in the number of single molecule, it also shows a direct transition from blue to red clusters and vice-versa (see, Fig. 7A). Overall, this suggests an increase in red-clusters as compared to blue clusters, indicating growth of HA clusters post, 24 hrs of transfection. In addition, we have carried out single molecule analysis to understand the HA clustering process in cellular system during influenza (type-A) viral transfection study (see, Fig. 7B1-B3). Here, a separate color-bar is used to represent time-dynamical changes for growing or diminishing clusters. Alongside the changes in these parameters (mean values) overtime are also shown in Fig. 7B4. The overall statistics is shown in Fig. 7B5 that indicate that fully-formed clusters have retained while a small section of clusters have diminished. However, a significant number of new clusters have formed over the time of investigation (∼60 *min*), post 24 hrs of transfection. The statistics suggests a dynamic situation where a significant number of new HA clusters have formed over time, and more importantly a large number of formed clusters have retained. This signifies a consistent formation of HA clusters and its growth over time, post 24 hrs of transfection.

**FIG. 7:**
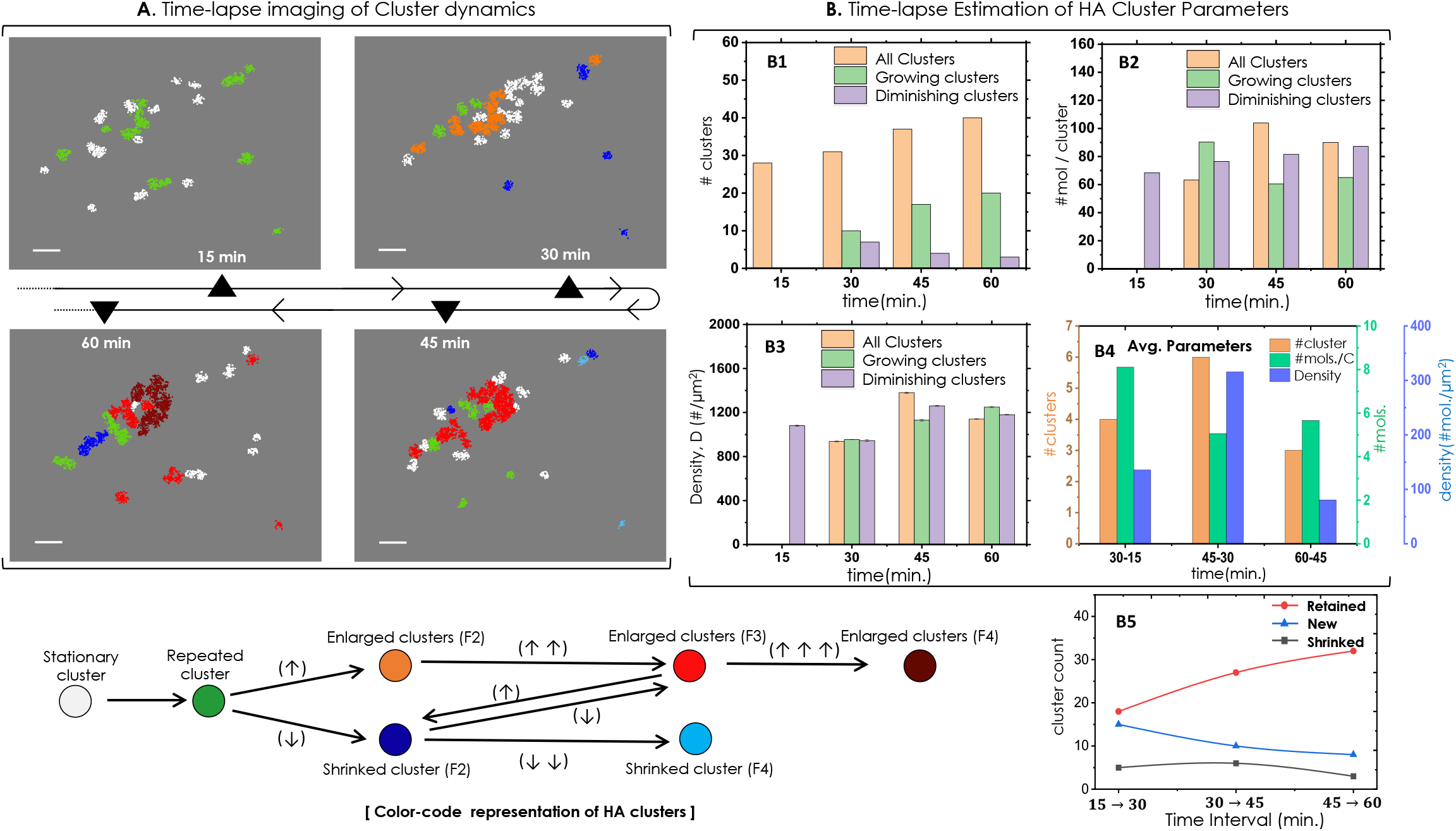
Tracking Single Molecule Clusters & Cluster Statistics: [A] Time lapse imaging of HA molecules and their clusters tracked on a time scale of 15 minutes and over an observation time of 1 hr. The clusters are colored following color code representation. [B1-B4] Estimation of dynamically changing cluster_-_ parameters (total number of clusters including clusters with increasing/decreasing HA molecules, the number of molecules in each cluster,the changes in density of the clusters), and the average parameters at varying time-points, post transfection. [B5] Alongside, the number of retained clusters, shrinked clusters,and newly formed are also shown. Below, color code representation for the state of dynamic clusters is also shown. Stationary clusters are marked by white color, while repeated clusters are marked by green color. The clusters that are growing (measured in terms of cluster size) are indicated by orange, red and brown color, with intermediate arrows indicate continuous growth. The clusters that are shrinking in size are indicated by dark and light color with arrows pointing down. The cluster growth and decline levels are indicated by number of arrows. Scalebar = 1*µm*

The experiments on fixed and live cells clearly indicate the advantage of event-based detection over frame-based detection employed in the existing SMLM techniques. With further development of event-based detection, single molecule imaging techniques are likely to play key roles in terms of high spatio-temporal super-resolution and low-cost of ownership.

## III. DISCUSSION

Single-molecule super-resolution microscopy is taking biology by storm, revealing events that were once thought impossible. Several studies have been reported over the last decade that demonstrate the capability of SMLM to reveal biological processes with single-molecule precision. Much credit goes to the development of new detectors that have accelerated SMLM imaging [52]. However, existing SMLM has poor temporal resolution, and the need for spatial resolution in sub-10 nm regime has increased over time. In this respect, event-based super-resolution microscopy is likely to play a big role, which is predominantly due to its ability to detect events (temporal change in the number of photons), and negligible readout time. With the availability of bright fluorophores and high quantum efficiency of the event detector, the technique is expected to push the limits of spatiotemporal particle resolution to < 1 *par* (where 1 *par* = 10^−11^ *ms*).

The fact that the event camera is sensitive to changes in intensity gives it the additional capability to record polarity (positive / negative changes in intensity). Unlike the existing SMLM that employ EMCCD/sCMOS detectors, event detector can continuously take data with negligible read-time. This is incredible since no signal / photon is lost during the entire period of investigation. However, even cameras have latencies, refractory period and quantization noise. For the present study, *eventSMLM* is first demonstrated by simply switching on/off the photon stream of a light source and subsequently using Cy3 blinking (see Fig. 2). Interestingly, Cy3 imaging has revealed non-zero events (change in photons) within the blinking period comprising both positive / negative events that corresponds to a temporal increase / decrease in recorded photons. This exposes the characteristic nature of photon emission from a single molecule within the blinking period.

Since event camera does not directly reveal the number of photons rather gives the change in the number of photons, the use of Thomson’s formula may not be necessary [50]. To deal with this, the localization precision is calculated geometrically as discussed in **Supplementary Note 4**. The method determines centroids from multiple realizations / *tPSFs* of single-molecule collected within the blinking period. From this set of centroids, a circle is fit, giving its center and radius. Subsequently, a Gaussian function is fitted from which the centroid and FWHM are determined. This corresponds to the position and geometric localization precision of the molecule. The same is repeated for all the molecules and a super-resolution map is reconstructed. Unlike Thompson’s formula, geometric localization precision is purely data-driven. For example, a spatial resolution (*R* = 2.3*l*_*p*_) of ∼15 *nm*) along with a temporal resolution of ≤ 10 *msec* produced an image with a spatio-temporal resolution of < 15*p*. In case of fixed structures and bright molecules (such as Cy3), the spatio-temporal resolution can be upto 2.3*l*_*p*_ × *τ* = 2.3 × 4.581*nm* × 3*msec* ≈ 3 *par*. This is a multi-fold improvement over the existing SMLM techniques.

The proposed microscopy technique is employed to visualize the arrangement of mitochondrial membrane protein (Tom20) labeled with mEos (see Fig. 3). The results revealed ∼7-fold improvement in localization precision, apart from high temporal resolution. In addition, the technique is used to investigate HA clusters in transfected NIH3T3 cells with an impressive localization precision of 6.56 *nm* and a 6-fold improvement in temporal resolution (see Fig. 4). Altogether, an improvement in spatio-temporal resolution of > 17 is noted when compared to standard SMLM. The study over a range of collection time and its effect on the quality of the reconstructed image (with respect to HA clusters) indicate the robustness and consistency of the proposed microscopy technique for a wide range of studies on both fixed and live cells (see Fig. 5). The technique (*eventSMLM*) further allowed determination of biophysical parameters that are directly linked to cell physiology (see, Fig. 6 and **Supplementary Note 8**). Finally, the technique was tested on live cells (post 24 hrs of transfection) to visualize the formation of HA clusters over a period of time. For the first time, we could visualize the formation of HA clusters using event-based single-molecule detection. This is in addition to the estimation of dynamic biophysical parameters (# clusters, #*HA*/cluster, cluster density, and cluster-fraction) (see, Fig. 6). In addition, time-lapse imaging shows dynamical formation and disappearance of HA clusters, with an over-whelming increase in cluster fraction over time. Overall, the advantage of event-based SMLM is apparent considering the complexity of processes involved in cellular system.

Future developments are likely to see an increased use of event cameras for single-molecule imaging. Event cameras with better quantum efficiency are likely to improve the temporal resolution in micro/nano-second time scale, thereby further pushing the limit of spatio-temporal resolution to reveal biological processes in action.

## IV. MATERIALS AND METHODS

### A. Toggle ON/OFF event setup

To experimentally generate ON/OFF events, we build an optical system, as shown in the Fig. 1. Complete details of each component can be found in **Supplementary Note 2**. A continuous beam of 532*nm* approximately 1*mm* beam diameter is expanded 4*X* times by using a beam-expander system that comprises of two biconvex lenses of focal lengths, 2.5*mm* and 10*mm*. The expanded beam is spit into two paths using a beam splitter 50 : 50. Each split beam is focused on the detector by Olympus 10*X*, 0.3*NA* objective lens. One of the focused spots is projected on the event camera, and the other spot is projected in a frame-based camera (CCD/EMCCD). To toggle the 532nm continuous laser ON/OFF, we used an electronics chopper in the path of the beam. The toggling ON/OFF and camera capture is synchronized using a microcontroller (Arduino). Toggle ON/OFF time can be set in the script code.

### B. Optical setup of *eventSMLM*

The key optical component of the *eventSMLM* system along with other details can be found in **Supplementary Note 3**. Light beams from activation excitation lasers of wavelengths, 405 nm and 561nm are combined by a dichroic mirror, and the combined beam is coupled with IX81 Olympus inverted microscope (used in epifluorescence mode). Illumination and collection of the fluorescence were done with oil immersion Olympus 100x 1.3NA. The dichroic filter instated in the microscope helps to distinguish excitation and emission wavelengths. Distinguished fluorescence is directed to the detection subsystem that is capable of imaging two channels to facilitate simultaneous capture of the signal using an event camera and EMCCD. We used beam splitter to simultaneous capture signals in both the detectors. In the EM-CCD detection path, an additional magnification of 2X is introduced to satisfy the sampling criteria. A set of optical filters are used in the detection path to collect fluorescence signal. The present configuration results in an effective pixel size of 45nm and 85nm for the event camera and EMCCD, respectively.

### C. Data acquisition

The data acquisition begins with identifying transfected cells using a separate fluorescence arm (not shown). Subsequently, the cell is prepared for super-resolution imaging. For quality imaging, the lasers are optimized for stable blinking of the molecules by matching the intensities that control the activation and emission rates [3]. A laser power of 20mW and 6*µW* for 561nm and 405nm is used. All data presented in this manuscript is acquired simultaneously in both the detectors. In EMCCD, the data are acquired in a stack of .tif format with 30ms exposure. Event camera parameters are optimized and key imaging parameters such as, field of view(FOV), sensitivity, and polarity sensitivity are set for data collection. Typical event detector parameters used while recording data arebias diff: 69, bias diff off: 52, bias diff on: 97, bias fo n: 0, bias hpf: 0, bias pr: 131 and bias refr:45.

### D. Event based frame generation

Any pixel of the event camera records a stream of events (*x, y, t, p*) where, (*x, y*) is pixel coordinate, *t* is the time at which intensity change occurs, and *p* is the polarity of events. Positive polarity corresponds to positive change in intensity; the corresponding event is called positive event, whereas negative polarity corresponds to a negative change in intensity, and the corresponding event is called negative event. One can reconstruct frames from any event type, i.e, positive event frames, negative events only frames or including both positive and negative events frame. Images of 8-bit .tif format of 10ms,30ms,60ms and 100ms time binned events (both positive and negative) are generated. The code for generating from events is written in phyton script. Details on different event frame construction algorithms can be found in Ref. [54]. Details on working of event camera can ve found in **Supplementary Note 1**.

### E. Event based image reconstruction

A customizable Matlab script is written to reconstruct the superresolved image. Approach similar to conventional super-resolved image reconstruction are applied. Initially, event-based particle identification is carried out. This pre-processing step helps in reducing unwanted signals. The left-out particle-like spots are identified using a correlation approach. An ideal PSF (diffraction-limited Gaussian PSF) is correlated with the pre-processed image, and based on the threshold correlation value, particles are identified (see, **Supplementary Note 4**). Only the particles that repeat in subsequent frames (at-least 3 frames) or more are considered to be molecules (within the blinking period of a molecule). A circle is fit through the centroids for which centre and diameter is determined. A 2D Gaussian function is invoked to represent single molecule PSF and a known relation (*FWHM* = 1.18*R*) between the radius of the circle (of radius, *R*) and FWHM of the Gaussian (*FWHM*) is used. The centre and 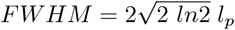 is used to estimate the position and localization precision (*l*_*p*_) of the single molecule see, **Supplementary Note 4**. Note that, the intensity of the molecules is proportional to the sum of events in each frame. Accordingly, we have the estimated information related to the molecule (its centre, localization precision, and intensity). Hence in the reconstructed super-resolved image, each detected molecule is represented by a Gaussian with its centre as the position, and the spot size as the localization precision. More details can be found in **Supplementary Note 4**.

### F. Goodness of Fit Test

To access the circle fitting through centroids, we used *R*^2^ goodness measure metric. This measure necessitates computing the residuals after fitting the circle. Residual sum of squares (*SS*_*res*_) measure how well the circle fits the data points whereas total sum of squares (*SS*_*tot*_) measures the variance of the data points around their mean. Goodness of fit is Calculated as, *R*^2^ = (1 − *SS*_*res*_)*/SS*_*tot*_. The *R*^2^ value gives you a normalized measure of how well the circle represents the data (centroid points), with 1 indicating a perfect fit. We found an average goodness of fitting value of 0.9 for *Cy*3 sample, indicating a good fit.

### G. Fluorescent proteins

To begin with the event-based detection of single molecules, we used the well-known fusion fluorescence protein mEosTom20. The plasmid is frequently used to visualize the Tom20 protein molecules, which are located at the mitochondrial outer membrane. The plasmid were purchased from Addgene, USA. The non-activated mEOSTOM20 can be visualized at 532nm upon blue 488nm excitation. After absorption of 405nm light, the absorption peak of molecules shift to 563nm and emission peak to 573nm.

Another fusion plasmid, Dendra2HA, is used to visualize the HA molecules in the plasma membrane of the cell after 24 hours post-transfection. HA is an influenza-type A viral protein. Event-based detection of HA molecules in fixed and live cells is demonstrated post-transient transfection.

### H. Cell culture and transfection

The study used mouse fibroblast NIH3T3 cell lines. 3T3 cells preserved in LN2 was thawn and cultured using standard protocols. After two passages, the cells were seeded with a count of approximately 10^5^ and cultured in a 35mm disc with coverslip for transfection. Post 24hr, the cells were used for transfection. Lipo P3000 transfection protocol is used to transfect cell with both the plasmids dendra2HA and mEOSTOM20. The concentration of plasmid used are 1078 *ng/µl* and 967 *ng/µl* for dendra2HA and mEosTom2O, respectively. After 24hr, cells were washed with PBS three times before fixing with 3.7% paraformaldehyde. Fixed cells are mounted with fluor save and air tighten by nail paint. For live imaging, 3T3 cells were grown in live imaging dish and same transfection protocol is adopted. Imaging is carried out in growth median after 24hrs of transfection.

## V. SUPPLEMENTARY NOTES

Supplementary Note 1: Event camera: working principle and calibration.

Supplementary Note 2: Toggle ON/OFF Event detection with event camera.

Supplementary Note 3: Complete optical setup of the eventSMLM system.

Supplementary Note 4: Computation of event based geometric localization of single molecules and image reconstruction.

Supplementary Note 5: Geometric Localization Precision at Varying Event Collection Time.

Supplementary Note 6: Super-resolving mitochondrial network (mEos-Tom20) in cellular system.

Supplementary Note 7: Event based super-resolution imaging of Dendra2-HA clusters in transfected cell.

Supplementary Note 8: FRC analysis for eventSMLM reconstructed images.

Supplementary Note 9: Estimation of critical biophysical parameters.

Supplementary Note 10. Event based imaging of cluster dynamics in a live cell.

## Contributions

PPM and JB conceived the idea. JB, AS, NP and PPM prepared the samples, and carried out the experiments. JB, AS, NP, VR, CST, PPM analyzed the data and prepared the figures. PPM wrote the paper by taking inputs from all the authors.

