## Supplementary Notes 1 - 10 for "Event-based Single Molecule Localization Microscopy (*eventSMLM*) for High Spatio-Temporal Super-resolution Imaging"

Jigmi Basumatary[1], Aravinth S[1], Neeraj Pant[1], Vignesh Ramanathan[3], Chetan S. Thakur[3] and Partha P. Mondal[1,2]
[1] Department of Instrumentation and Applied Physics, Indian Institute of Science, Bangalore 560012, INDIA
[2] Center for Cryogenic Technology, Indian Institute of Science, Bangalore 560012, INDIA
[3] Department of Electronics System Engineering, Indian Institute of Science, Bangalore 560012, INDIA

#### Supplementary Note 1-10

**Supplementary Note 1:** Event camera: working principle and calibration

**Supplementary Note 2:** Toggle ON/OFF Event detection with event camera.

**Supplementary Note 3:** Complete optical setup of the *eventSMLM* system.

**Supplementary Note 4:** Computation of event based geometric localization of single molecules and image reconstruction .

**Supplementary Note 5:** Geometric Localization Precision at Varying Event Collection Time

**Supplementary Note 6:** Super-resolving mitochondrial network (mEos-Tom20) in cellular system.

**Supplementary Note 7:** Event based super-resolution imaging of Dendra2-HA clusters in transfected cell.

**Supplementary Note 8:** FRC analysis for *eventSMLM* reconstructed images.

**Supplementary Note 9:** Estimation of critical biophysical parameters.

**Supplementary Note 10.** Event based imaging of cluster dynamics in a live cell

#### Supplementary Note 1

#### Event camera: working principle and calibration

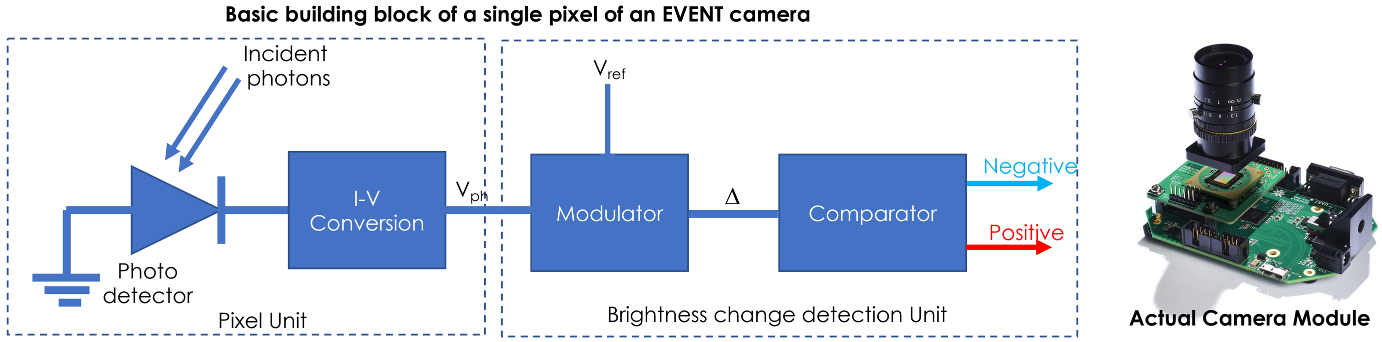


**Fig. S1-1. Basic Structure of Event Camera.** Incident photons are detected by a photo detector (photodiode), which converts the incident photons into electric current. The photon electric current is converted into voltage for further brightness change detection. The modulation unit calculates the difference between the reference voltage and the output voltage from the pixel unit. The comparator unit outputs a positive or negative event if the delta (Δ) exceeds the positive or negative threshold respectively. Alongside, the actual picture of commercially available event camera is also shown [1,2].

Broadly, the basic building blocks of event camera are the pixel unit and the detection unit. The pixel unit converts the incident photons into electric current, and the detection unit determines change in brightness (both positive and negative) (see, Fig. S2-1). Specifically, the camera sensor consist of rectangular array of pixels and each pixel works independently. Each pixel (light receiving unit) of the event camera is connected to electronics circuit (luminance change detection unit) in such a way that pixel sensor together with electronics circuit respond to optical intensity variation only. Since each pixel works independently, the functioning of event camera can be understood based on the operation of a single camera pixel. At any time t, a pixel convents incident light intensity I(t) into electronic response and store a reference level $I_{ref}$. The variation in light intensity is detected relative to $I_{ref}$ based on two thresholds: T- and T+. Threshold T- is used detect decrease in intensity and T+ is used to detect increased in intensity. The user can set the threshold values as per the desired application. In our experiment we used same value (11% of the reference value) for both the thresholds. Detection of the variation in light intensity is in logarithmic scale. This facilitates sensor to detect subtle difference in the low intensity range while it responds to a wide intensity difference in the high intensity range to prevent event saturation, allowing it to realise a wide dynamic range. Hence an event signal is triggered as soon as $I(t)/I_{ref} > T+$ (positive event) or $I(t)/I_{ref} < T-$ (negative event). Whenever event is triggered the sensor returns a signal containing pixel address, polarity of the event and time stamp of the event i.e, (x,y,p,t). After the event is triggered the $I_{ref}$ is updated with the value of intensity at the time of trigger. Hence the output of event camera is sparse list containing (x,y,p,t). Time resolution of the sensor is limited by refractory period which is of the order of microseconds [3]. During death time sensor cannot trigger event. For event based super-resolution imaging, the knowledge of position (x,y), polarity (p), and time of the event (t) is sufficient to super-resolve point / single molecule emitters. More details on event camera can be found in review paper [3,4].

**Supplementary Note 2**

#### Toggle ON/OFF event detection with event camera

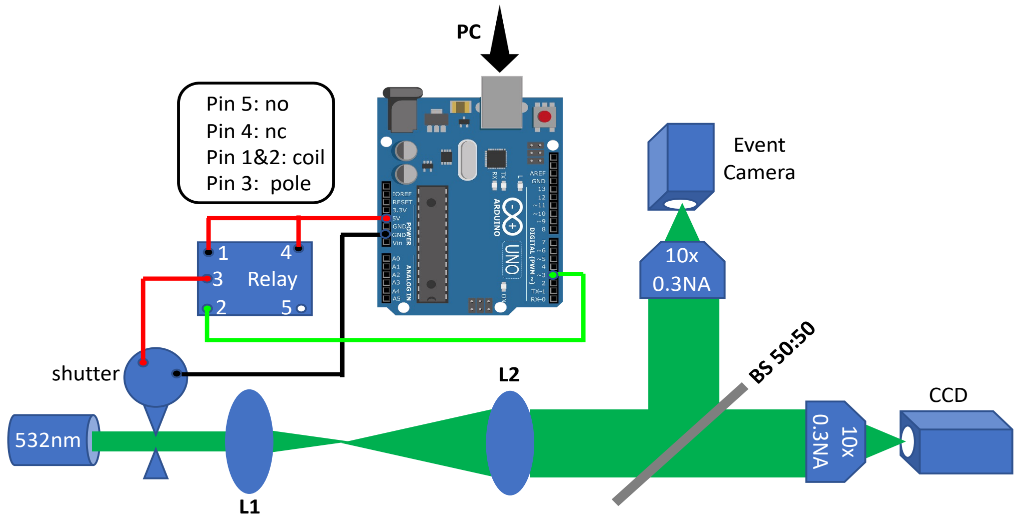


**Fig. S2-1. Event Detection.** ON/OFF events of 532nm continuous laser. Beam shutter is motorised in which voltage applied to is controlled via relay. Relay is operating at normally closed (nc) mode. The frequency of relay switching is programmed with Arduino.

To facilitate optimal working and calibration of event camera for event-based detection, we built a simple optical setup for simultaneous detection as shown in FigS2-1. The beam of 532nm laser is expanded 4 times by a beam-expander (comprises of lenses, L1 and L2), and split into two paths by a 50:50 beam splitter. Further each beam is focus to diffraction limited spot by a 10X, 0.3 NA objective lens. The key components in the system are:

1. 532nm Laser source: Laserglow, LCS-0532-TDD-00100-05, Avg power=207.2 mW).
2. Shutter: Thorlabs, SH1/M, 10ms response time.
3. Lens L1 and L2: f1=2.5cm and f2=10cm, Thorlabs
4. CCD: point grey, CMLN-13S2M-CS, pixel size 3.75 µm
5. Event camera: PROPHESEE, EVKV2_gen 4.1, pixel size 4.5 µm
6. Objective Lens: O1 and O2 , Olympus 0.3 NA

A chopper is used in the path of the beam to control ON / OFF during the experiment to mimic change in the intensity. The chopper is motorised and it's on/off frequency is programmed via relay (5V, EC-4504, electronicscomp.com) through Arduino microcontroller (UNO R3). The complete connection setup is shown in Fig. S2-1. The cameras (Event and CCD) are not synchronised but data is recorded simultaneously.

#### Complete optical setup of the *eventSMLM* system

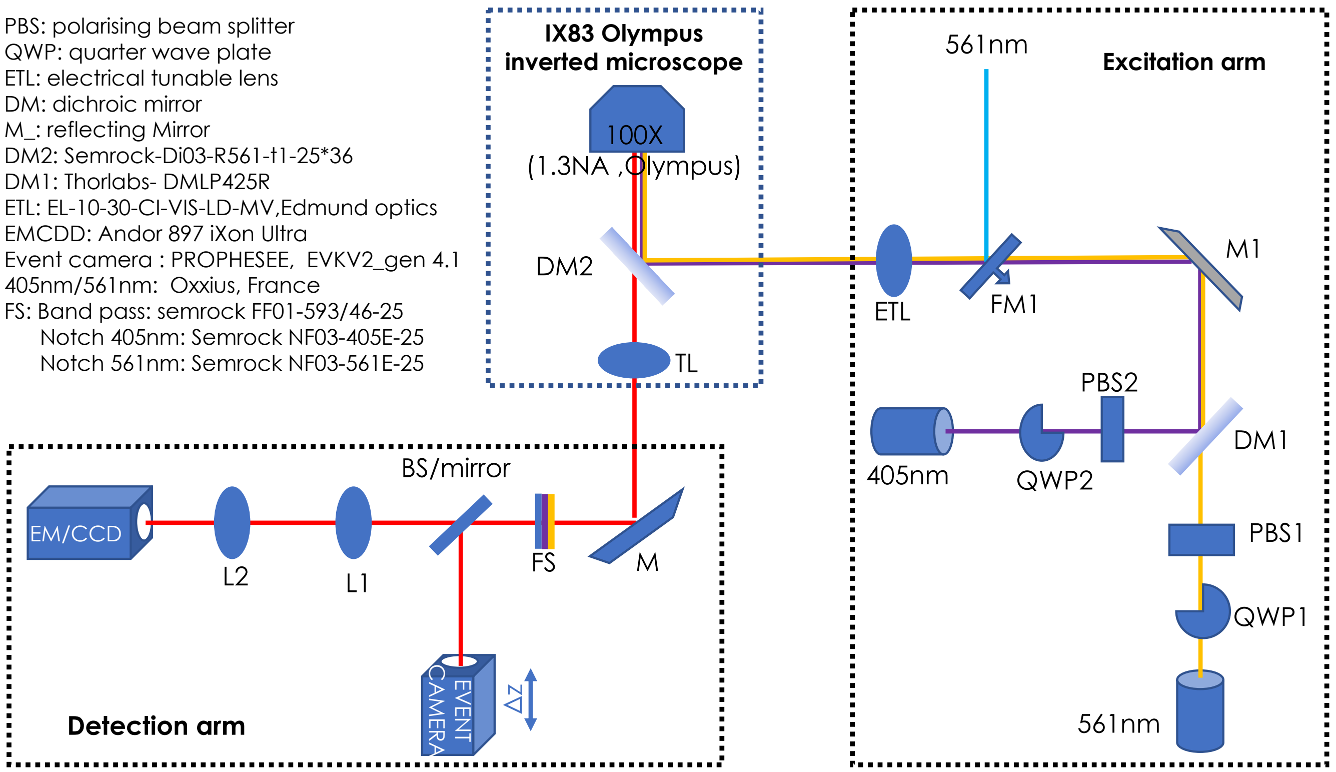


**Fig. S3-1. Schematic diagram of *eventSMLM* system.** A complete schematic of developed *eventSMLM* system shows the optical arrangement in the actual experiments along with the key optical component (with part number). Three major sections are mentioned: excitation arm, detection arm and the inverted microscope.

An epifluorescence mode SMLM system is used to realize *eventSMLM* system as shown in Fig. S3-1. The complete setup with key optical components along with part number is also shown in Fig. S3-1. The excitation arm consists of activation laser and readout laser which are combined with a dichroic mirror. In each beam path, intensity control unit made up of PBS and QWP are installed. Combined beam is directed to the high NA objective lens of the inverted microscope. ETL lens in the excitation arm is used to set the appropriate field of view. The fluorescence signal from the single molecules is separated by the dichroic mirror DM2 and is focused by tube lens to the primary image plane. The detection arm is customised to image signal simultaneously into two channels by the beam-splitter BS. In place of BS, mirror is used to image fluorescence with event camera only. Event camera captures the primary images, while EMMCD captures the magnified version of the primary image. Hence the event camera detection has effective pixel size of 45nm whereas EMCCD has a total magnification of 266X, giving an effective pixel size of 62.02nm. The setup is used to visualize single molecules (mEos-Tom20) distribution on the mitochondrial network. In addition, the technique is further employed to study understand HA clustering in transfected NIH3T3 cells.

#### Computation of event based geometric localization of single molecules and image reconstruction

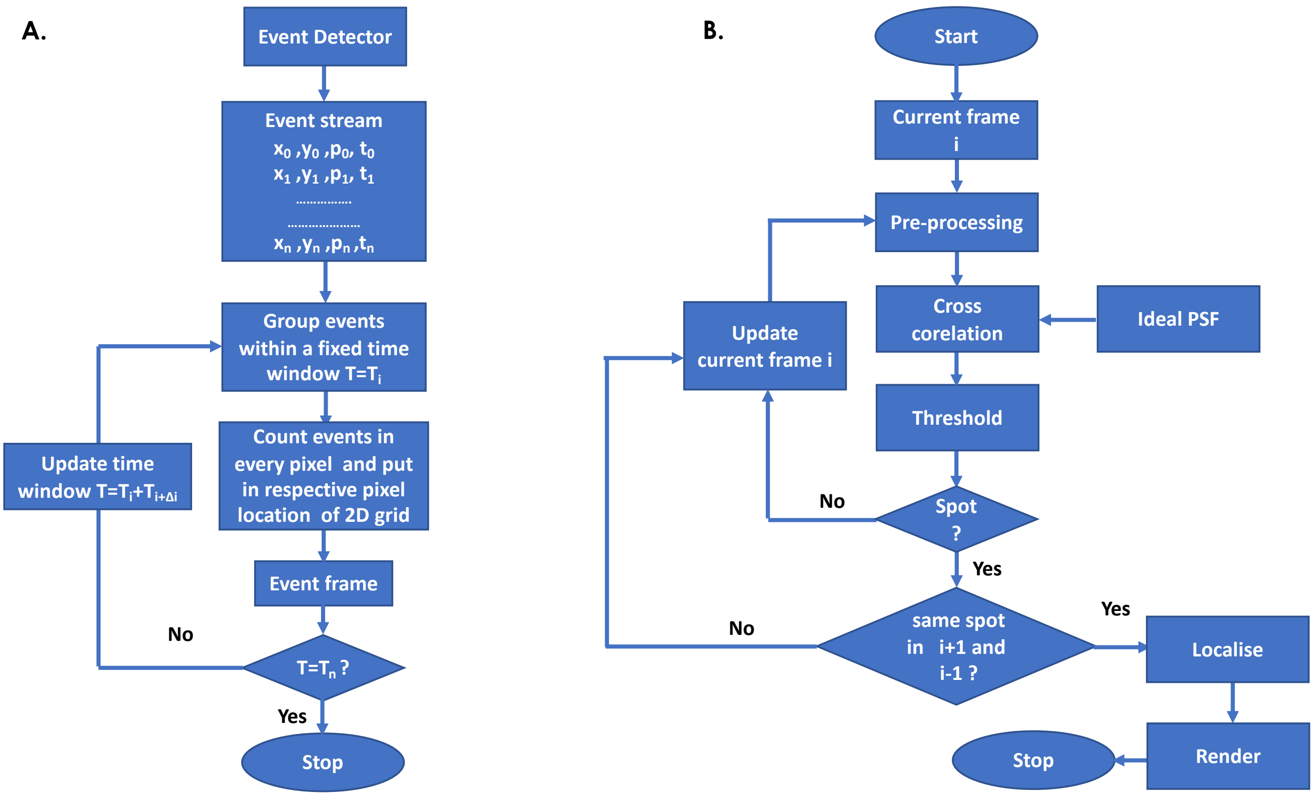


**Fig. S4-1. Flow chart for computation of localization precision and image reconstruction.** (A) frame reconstruction from stream of events (B) Steps involved in event based single molecule reconstruction. Briefly event-based frames are pre-processed to remove unwanted noise. The frames are correlated with ideal PSF, and a suitable threshold is used to extract the spots from frames. Algorithm also look for the events of the same molecules in the subsequent frames.

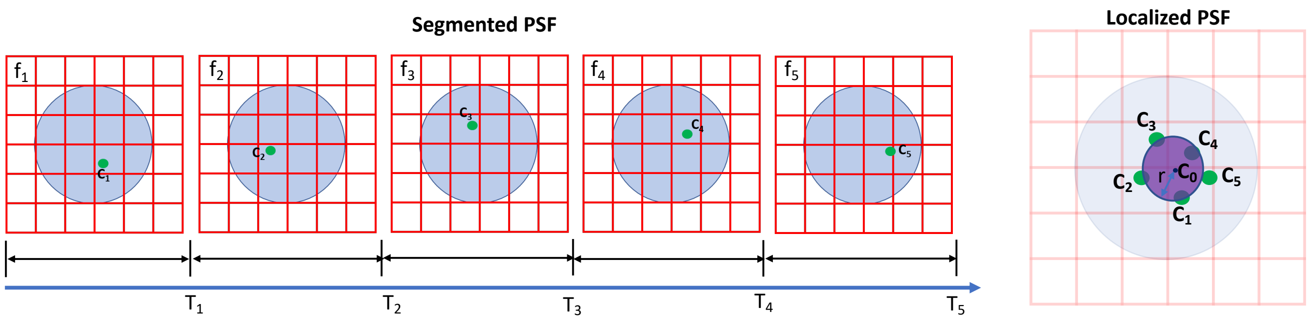


**Fig. S4-2. Computation of Geometric Localization Precision.** During the ON time of the single molecule T=T1+T2+T3+T4+T5 multiple event frames f1, f2, f3, f4, f5 which are also called segmented PSF, are generated by collecting events at a fixed event collection time. In each frame intensity weighted centroid are calculated C1, c2, c3, c4, c5. A circle is fitted passing through these centroids and the parameters (center, Co and radius, r) are determined. Localisation precision is defined by the size of the circle.

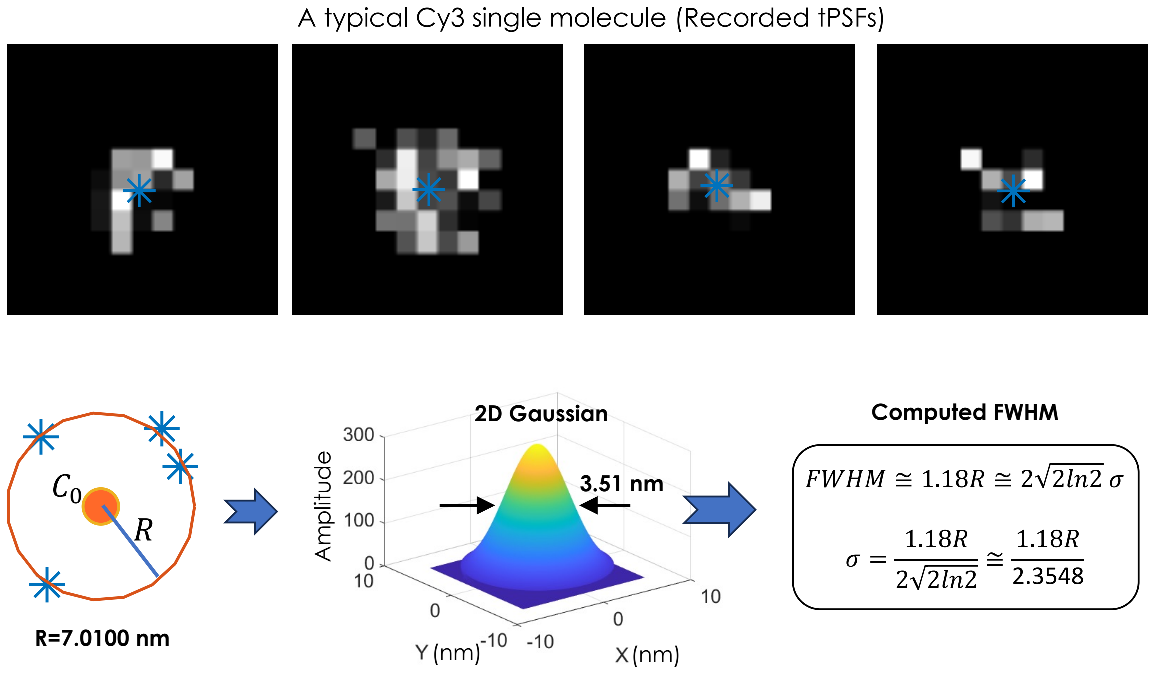


**Fig. S4-3. Computation of Localization Precision.** Recorded PSFs of a typical single Cy3 molecule for a collection time of 100 ms. The corresponding fit-circle followed by a 2D Gaussian function determine its location and geometric localization precision.

To reconstruct single molecule using *eventSMLM*, all the detected events within the blinking time of a molecule are recorded. From event detector data (x,y,p,t), we reconstruct event based frame by collecting events in a fixed window of time (event collection time). Gray level of the spots in the reconstructed frame is equal to the number of events detected in the collection time. Each pixel of event detector records stream of events with *p* as the polarity of events. Positive polarity corresponds to positive change in intensity, whereas negative polarity corresponds to negative change in intensity. The flow chart for event based reconstruction in shown in Fig. S4-1.

Briefly, the molecule events are segmented into multiple frames by time binning as shown in Figure S4-2. In each frame centre of mass of the spot is estimated. We defined the position of molecule by centre Co(a, b) of the fitted circle through the centroid of each frame, and localisation precision by the FWHM of the fitted Gaussian. This gives position of the molecule and its approximate size (localization precision). As an example, the process is demonstrated for a typical Cy3 molecule for which PSFs are collected every 100 ms (see, Fig. S4-3). Each molecule is rendered as a 2D Gaussian spot having width equals to calculated localisation precession and intensity A is the sum of all the events (both positive and negative). Circle fitting through centroids of segmented PSF can be understood easily. Suppose we have collection of $n \geq 3$ centroid points in 2D-space labelled as $C_1(x_1, y_1)$, $C_2(x_2, y_2)$, . . . , $C_n$ $(x_n, y_n)$. With well-known equation of circle described by, $(x - a)^2 + (y - b)^2 = r^2$ , our objective is to estimate centre $C_0(a, b)$ and radius r for the best fitting circle. Reasonable measure of the fit of the circle, $(x - a)^2 + (y - b)^2 = r^2$ to the points $C_1$ , $C_2$ , . . . , $C_n$ is given by summing the squares of the distances from the points to the circle[1]. This measure minimizes the sum of squared radial deviations and is given by,

$$SS(a,b,r) = \sum_{i=1}^{n} \left(r - \sqrt{(x_i - a)^2 - (y_i - b)^2}\right)^2 \qquad (1)$$

where, ***a, b***, and ***r*** are chosen such that SS is minimized.

Differentiation of SS yields,

$$\frac{\partial ss}{\partial r} = -2\sum_{i=1}^{n} \sqrt{(x_i - a)^2 - (y_i - b)^2} + 2nr \qquad (2)$$

$$\frac{\partial ss}{\partial a} = 2r\sum_{i=1}^{n} \frac{x_i - a}{\sqrt{(x_i - a)^2 - (y_i - b)^2}} - 2nx + 2na \qquad (3)$$

$$\frac{\partial SS}{\partial b} = 2r\sum_{i=1}^{n}\frac{x_i - b}{\sqrt{(x_i-a)^2-(y_i-b)^2}} + 2ny + 2nb \qquad (4)$$

This will give the fitting parameters, *r, a, b* from which the centre, $C_0(a,b)$ and radius, *r* are determined. Subsequently, a Gaussian representation is used to render localised PSF, with $C_0(a,b)$ as the centroid, *lp* as the localization precision and $I_0$ as the peak intensity (sum of all events),

$$PSF_{loc}(a,b) = I_0\, e^{-\frac{(x-a)^2+(y-b)^2}{2l_p{}^2}} \qquad (5)$$

Next, we establish the relation between the radius *r* and *lp*. In general, a Gaussian PSF as a function of the radial distance $r$ from the center is given by [2,3],

$$I(r) = I_0 e^{-\frac{2r^2}{R^2}} \qquad (6)$$

where, $I_0$ is the maximum intensity at the center of the beam ($r = 0$), $R$ is the radius which is defined as the distance at which the intensity drops to $\frac{I_0}{e^2}$ .

Here, FWHM is the distance between the two points where the intensity falls to half of the maximum value. So, setting the intensity $I(r)$ equal to half of the peak intensity $I_0$ gives,

$$\mathrm{I(FWHM/2)} = \frac{I_0}{2}$$

$$I_0 e^{-2\frac{(\frac{FWHM}{2})^2}{R^2}} = \frac{I_0}{2}$$

Solving we have,

$$FWHM \cong 1.177\, R \cong 1.18\, \mathrm{R} \qquad (7)$$

Noting that the localization precision *lp* is related to the FWHM of the Gaussian i.e, $FWHM = 2\sqrt{2\, ln2}\; l_p$, we get,

$$1.18\, R = 2\sqrt{2\, ln2}\; l_p$$

$$\Rightarrow\; l_p \approx \frac{1.18\, R}{2.355} \qquad (8)$$

The above equation is used to calculate the localization precision of the single molecules.

[2] Book: Optics, by Eugene Hecht, Chapter. 13, 4th edition, Pearson, 2008.

[3]https://support.zemax.com/hc/en-us/articles/1500005488161-How-to-convert-FWHM-measurements-to-1-e-2-halfwidths

### Supplementary Note 5

#### Geometric Localization Precision at Varying Event Collection Time

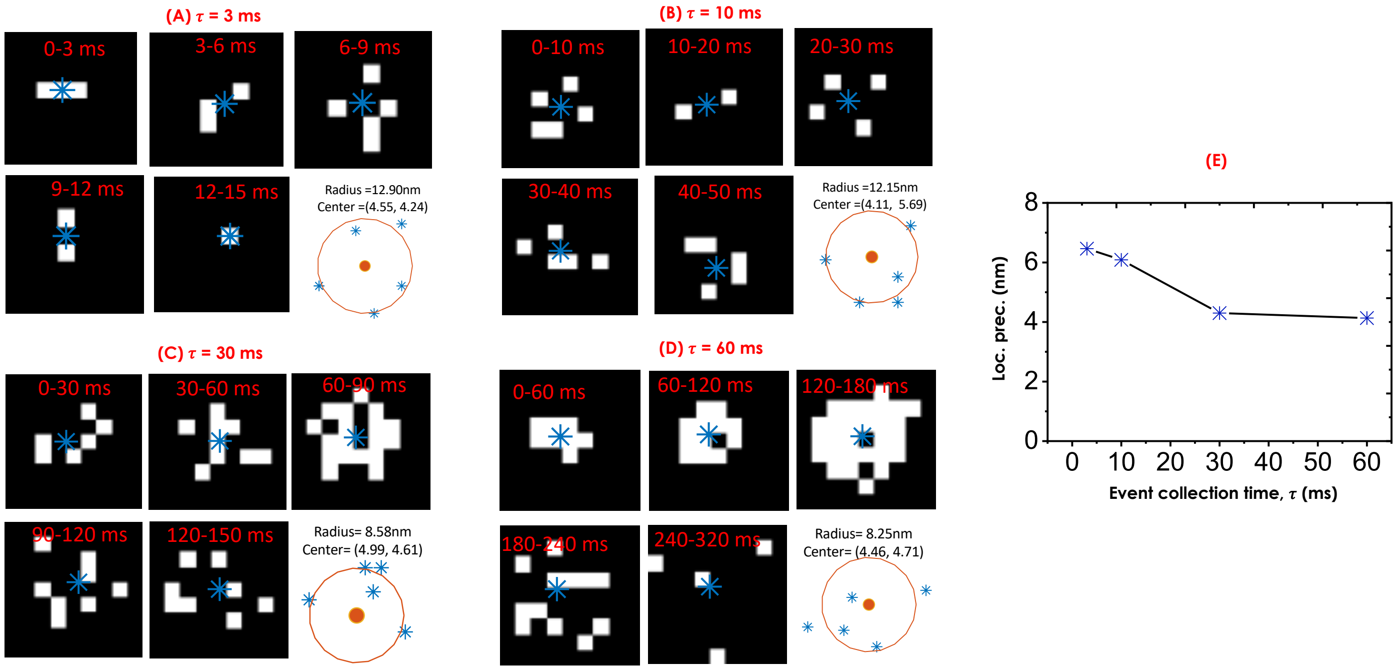


**Fig. S5-1. Geometric localization precision :** Evaluation of localization precision for multiple realization of PSF of Cy3 molecule at varying event collection times (3 ms, 10 ms, 30 ms). The corresponding centroid (center) and localization radius are also shown. Green asterisk represents the centroid of localization circle. (D) Plot of localization precision versus event collection time, showing an expected reduction in localization precision with large collection time (τ).

To understand the effect of event collection time on the geometric localization precision, single molecule experiments are performed on Cy3 molecules. For each case, 5 realization of the PSFs are taken and the corresponding centres, Co and geometric localization precisions are determined. Fig. S5-1 shows three selected cases where the event collection times are varied from low (3 ms) to high (30 ms). Specifically, 5 different PSFs are observed for each collection time and their centre and radius are computed using eqn.(1-5) (see, Supplementary Note 4). Although the PSFs are sparse (which is largely due to less photons per pixel, camera latency and short accumulation time), the centroids are close enough to give an approximate location of the molecule (geometric localization precision). The corresponding plot (Fig. S5-1D) shows a decrease in the localization precision with an increase in event collection time. This is understandable since large collection time produces more photons, thereby increasing the accuracy of localization precision. This overall improves spatio-temporal resolution of *eventSMLM* technique with the smallest value of, $l_p \times \tau = 3ms \times 12.90\, nm = 38.7\, nm.msec$.

#### Supplementary Note 6

#### Super-resolving mitochondrial network (mEos-Tom20) in a cellular system

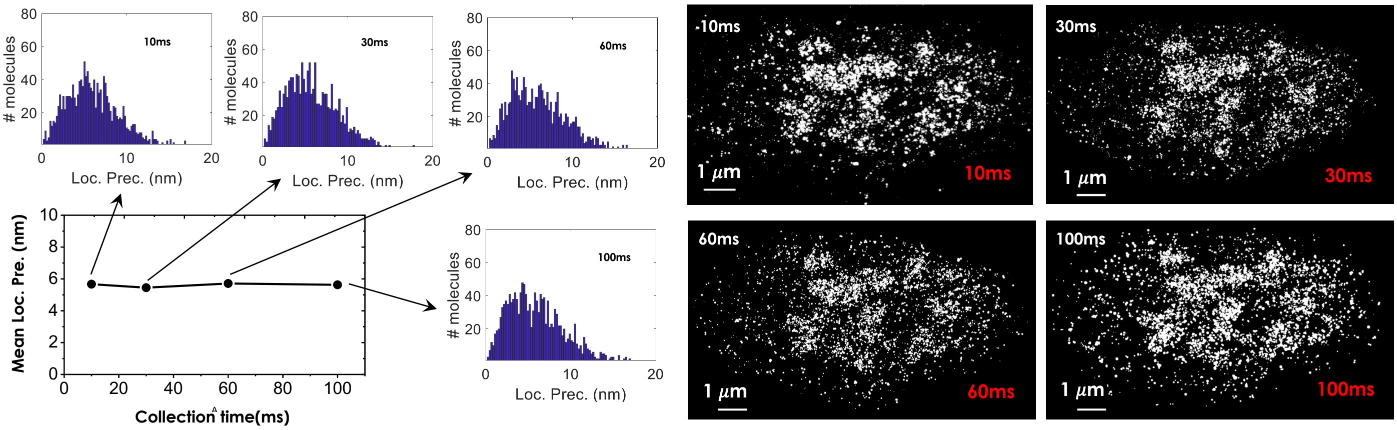


**Fig. S6-1. Reconstructing mitochondrial network.** [A] Event-based image reconstruction at varying event collection time. [B] The variation of localisation precision with collection time.

The *eventSMLM* reconstructed images of mEos-Tom20 molecules in a transfected NIH3T3 cell is shown in Fig. S6-1. It is evident that location of Tom20 accumulates and its key features are well-preserved, although the effect of small statistics is clearly visible at an event collection time of 3ms. The specimen is prepared following the protocol discussed in the main manuscript, section IV.G. The event based single molecule PSFs are recorded post 24hrs of transfection, and an approximate 5000 images are collected. Imaging is carried out by sending 100% florescence to event detector (using mirrors, see optical diagram in Supplementary Note 3). Alongside mean localization plot and histogram of localization precision (see, insets in Fig. S6-1A) at varying collection times (10-100 ms) is also shown. As expected, the localization precision average remains same.

#### Supplementary Note 7

Event based super-resolution imaging of Dendra2-HA clusters in transfected cell

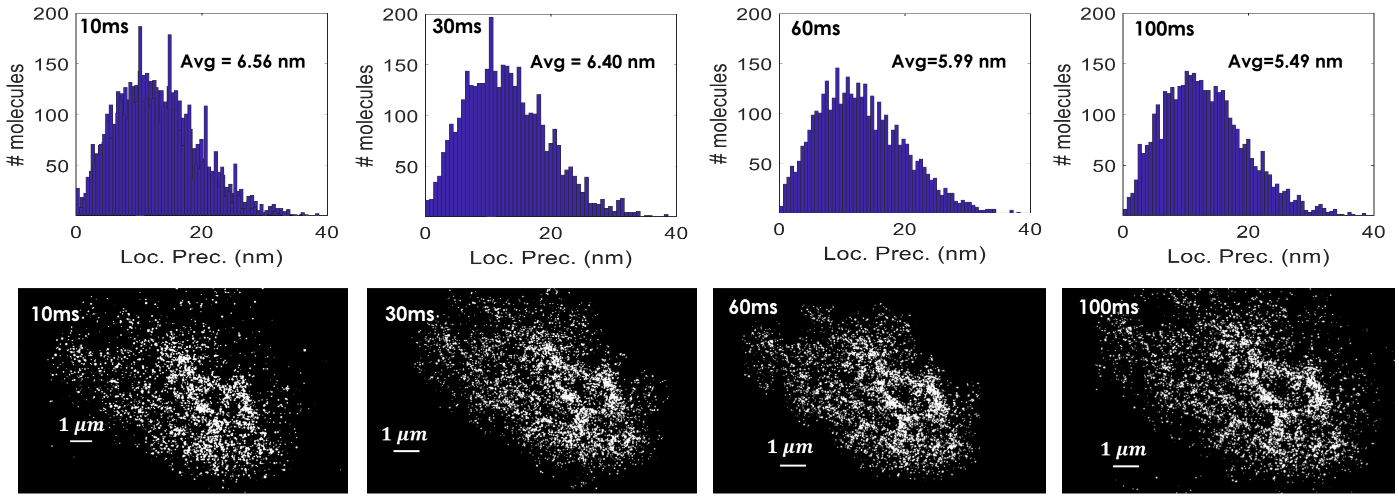


**Fig. S7-1. Super-resolving HA clusters.** Event based single molecule image of Dendra2-HA clusters in a transfected cell for varying event collection time. The corresponding event-based geometric localisation precision histogram is also shown.

The *eventSMLM* reconstructed images of Dendra2HA transfected cell. Single molecule event are collected post 24 hrs of transfection. Fig. S7-1A shows the images of HA clusters in a cell at varying collection times (10-100 ms). Again better localization precision is obtained at high collection time which is due to better photon statistics for individual molecule. The average mean resolution is ~ 12.91 nm for *eventSMLM* reconstructed images which is approximately 3.5 times lower than standard SMLM reconstructed images. Another distinct feature to note is the arrangement of HA clusters which appears better resolved at large collection time (100 ms) as compared to small collection time (10 ms).

**Supplementary Note 8**

FRC analysis for *eventSMLM* reconstructed images

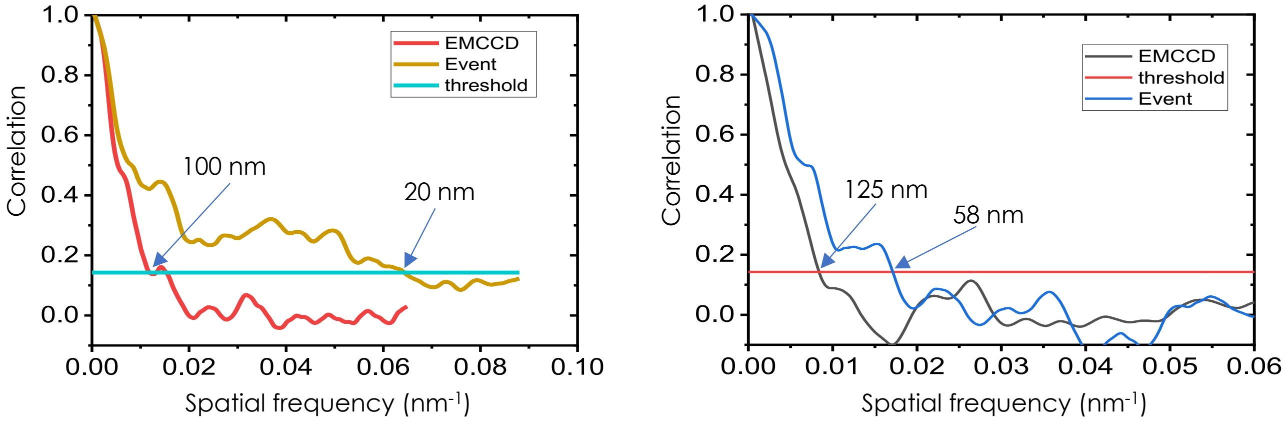


**Fig. S8-1. FRC Analysis.** FRC analysis of event based SMLM images of, [A] mEosTom20, and [B] Dendra2HA. At a standard cut-off of 1/7, resolution of mEOSTom20 and HA clusters are found to be ~20nm and ~58nm respectively.

Fourier Ring Correlation (FRC) is a method for calculating the correlation between two images as a function of spatial frequency. It can be viewed as a measure of the resolution of an image when applied to a pair of super-resolved images produced by separating the list of localization coordinates into two sub-samples. The two sub-images are correlated by multiplying their Fourier transformations Fx and Fy. The Fxy is calculated by summing over concentric rings and normalized by the total intensities in each ring in Fourier space i.e,

$$F_{xy} = \frac{\sum_{r \in r_i} F_x(r) F_y(r)^*}{\sqrt{\sum_{r \in r_i} {F_x}^2(r) {F_y}^2(r)^*}} \quad (1)$$

The signal at a distance $r_i$ from the center of the Fourier transformed images corresponds to the spatial frequency, $f_i = n/x_i$ , $n_i$ is the number of frequency bins, or pixels in the image. The spatial frequency where the FRC falls below a value of $^1/_7$ is defined as cut-off frequency and interpreted as resolution of the full image. The resolution analysis using FRC approach is calculated using Fiji as shown in Fig. S8-1(A,B) for mEos-Tom20 and Dendra2-HA. The *eventSMLM* reconstructed images shows an impressive 5 fold improvement for mEos-Tom20 and >2 fold improvement for Dendra2-HA. This is impressive considering the fact that event based reconstruction involves low-photon statistics which is due to change based nature of event camera and a relatively low quantum efficiency of the detector.

#### Estimation of critical biophysical parameters

| HA Cluster Parameter | Event detector (90%) | | | | EMCCD(10%) |
|---|---|---|---|---|---|
| | Event collection time | | | | Exposure time |
| | 10ms | 30ms | 60ms | 100ms | 60ms |
| Total area(μm²) | 5.05 | 5.34 | 5.13 | 5.21 | 4.17 |
| #mols. | 2516 | 2737 | 2661 | 2604 | 2650 |
| Density (#mols. / μm²) | 477.23 | 487.93 | 479.88 | 486.47 | 635.49 |
| Cluster fraction(%) | 11.09 | 11.73 | 11.26 | 11.44 | |

**Table T1. A comparison of** biophysical parameters (cluster area, #molecules / cluster, density and cluster fraction) associated with the HA clusters in a transfected NIH3T3 cell between event detector and EMCCD (exposure time 30 ms).

The proposed *eventSMLM* technique allows computation of biophysical parameters based on the event detection (single molecule blinking). In the cell transfection study, HA molecules are found to form cluster post 24 hrs of transfection. We have used a well-known DBSCAN clustering algorithm to identify the clusters. Specifically, single molecules closer than 30 nm is used as a cut-off for identifying molecules belonging to a particular cluster. In addition, a minimum number of points greater than 200 is considered for identifying a valid cluster. The clusters are identified at different event collection times (10-100 ms) and compared with standard SMLM. The corresponding biophysical parameters are tabulated in table T1. The parameters indicate near consistency of cluster parameters across the event collection times. This indicates that the statistics related to HA clusters are preserved at low collection times as well, justifying the use of small collection times for better spatio-temporal resolution. This further states that *eventSMLM* is beneficial for imaging dynamic events (both single molecule and its collective motion) in cellular system.

#### Supplementary Note 10

Event based time-lapse imaging of HA cluster in a live cell

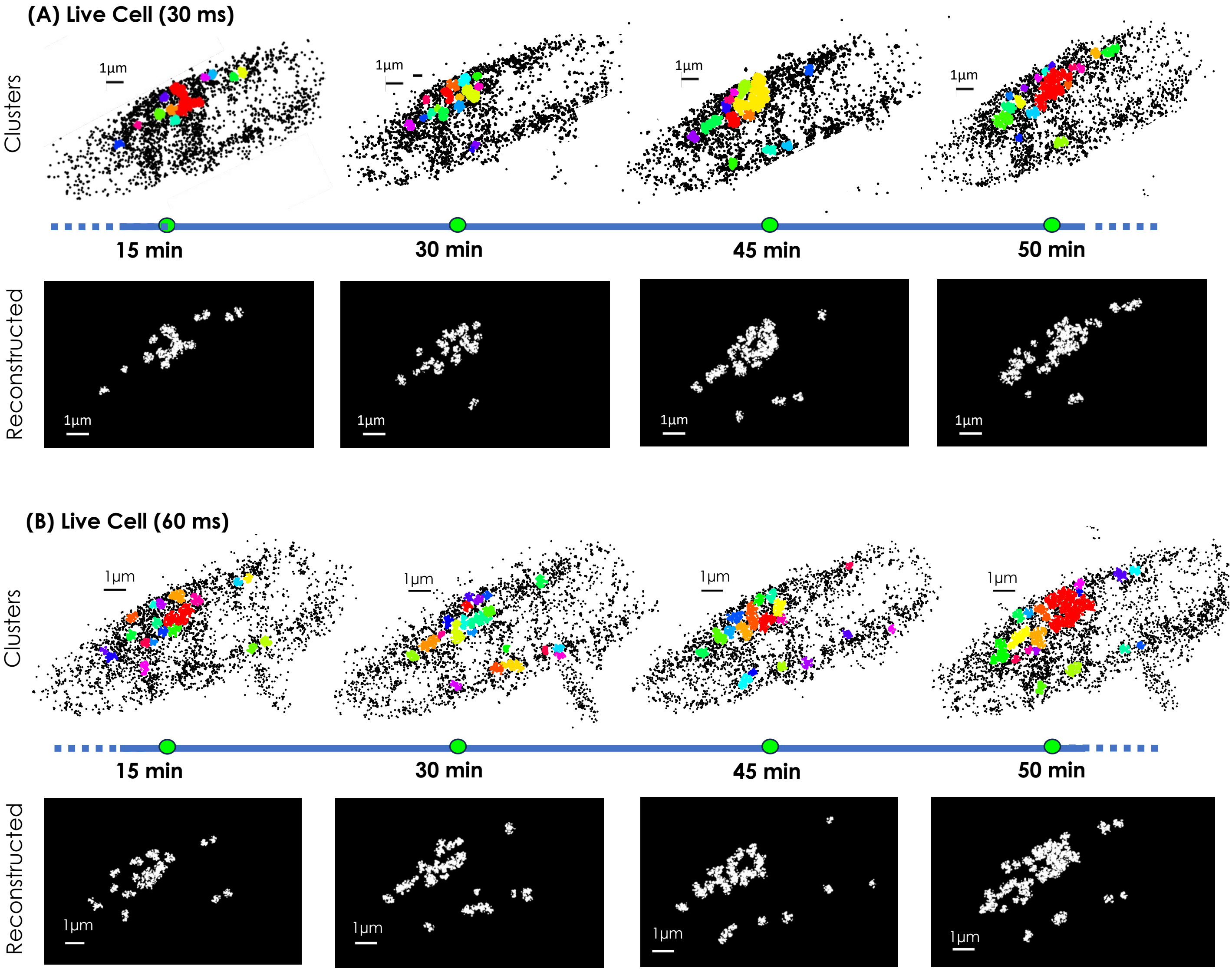


**Fig. S10-1. Time-lapse imaging of HA clusters over time**. Event based HA cluster dynamics in NIH3T3 cell with an event collection time of, [A] 30 ms, and [B] 60ms.

With the availability of *eventSMLM* and its high spatio-temporal resolution, we employed it to visualize and understand the dynamics of HA clusters in a live cell during early stages of viral infection as shown in Fig. S10-1. To begin with, NIH3T3 cell was transfected with Dendra2HA plasmid using Lipo-P3000 protocol. We recorded single molecule blinking events continuously for 60 minutes. The data is collected every 15 minutes and the corresponding super-resolved images are reconstructed. DBSCAN clustering algorithm is employed to identify HA clusters. Fig. S10(A,B) shows cluster dynamics enabled by event based detected with event collection times of 30ms and 60ms. The formation of clusters and its increase over an observation time of 60 minutes is quite evident. This clearly indicates the rate of HA clustering and its dynamics (migration/formation/dissolution) during Dengue viral infection.
